# Dynamin proline-rich domain isoform abundance, not identity, determines synaptic vesicle recycling efficiency in *Drosophila*

**DOI:** 10.64898/2026.08.24.746809

**Authors:** Anne M. Silveira, Kevin M. De Léon Gonzalez, Amy L. Scalera, Laura J. Westhoff, Kathy A. Roytman, Steven J. Del Signore, Bruce L. Goode, Avital A. Rodal

## Abstract

During neurotransmission, synaptic vesicle exocytosis adds membrane and proteins to the cell surface. To sustain further release, this material must be retrieved, via several distinct endocytic modes matched to the level of exocytosis. The GTPase dynamin plays a central role in endocytosis, but it has remained unclear which endocytic modes it supports. In mammals, distinct dynamin gene products with different proline-rich domains (PRDs) are proposed to mediate particular modes of endocytosis; however, the function of each PRD isoform has not been tested in an organism. *Drosophila* dynamin is encoded by one gene (*shibire*) that produces long and short PRD isoforms (Shi-L and Shi-S), which differ by a 48 amino acid C-terminal extension. Using isoform-specific knockin and knockdown tools, we found that loss of the more abundant Shi-S isoform disrupted bulk endocytosis and vesicle reformation under high exocytic demand, reduced evoked transmission at moderate levels of activity, and enhanced spontaneous release at rest. These functions did not depend on the PRD extension, as either isoform could rescue these phenotypes when re-expressed. Our results indicate that dynamin contributes to vesicle recycling across multiple endocytic retrieval modes and that PRD specialization is not required for these functions.

## Introduction

Synaptic vesicle (SV) exocytosis rapidly adds membrane and proteins to the neuronal surface. Subsequent recycling of these components by endocytosis is required to maintain membrane size and tension, clear release sites, and replenish SV pools (Kononenko and Haucke, 2015; Soykan et al., 2016). Multiple endocytic pathways contribute to SV recycling, including clathrin-mediated, ultrafast, kiss and run, and bulk endocytosis (Chanaday et al., 2019; Gan and Watanabe, 2018; Saheki and De Camilli, 2012; Watanabe, 2025). These pathways operate on different timescales and in response to different regimes of neuronal activity, but nonetheless rely on a common set of endocytic proteins (Azarnia Tehran and Maritzen, 2022; Dittman and Ryan, 2009). Whether specific proteins have unique functions and regulatory inputs in each of these endocytic pathways remains a central question in synaptic cell biology.

The large GTPase dynamin is a key endocytic protein involved in SV recycling. Dynamin contains an N-terminal GTPase domain, a stalk domain, a pleckstrin homology (PH) domain, a GTPase effector domain (GED), and a C-terminal proline-rich domain (PRD). Dynamin is best known for oligomerizing around the neck of invaginated membrane pits and using mechano-chemical energy released by GTP hydrolysis to drive membrane scission (Mettlen et al., 2009; Antonny et al., 2016) (Kokotos and Cousin, 2015).

However, dynamin has additional documented functions in membrane curvature sensing (Liu et al., 2011; Roux et al., 2010), actin regulation (Grassart et al., 2014; Gu et al., 2010; Taylor et al., 2012; Zhang et al., 2020), fusion pore dynamics (Wu and Chan, 2024), and scaffolding or regulatory interactions with SH3 domain-containing endocytic proteins (Meinecke et al., 2013; Rosendale et al., 2019; Sundborger and Hinshaw, 2014). These diverse activities may allow dynamin to function in different modes of SV endocytosis (Prichard et al., 2021), including clathrin-mediated endocytosis (Ferguson et al., 2007; Raimondi et al., 2011), ultrafast endocytosis (Imoto et al., 2022; Imoto et al., 2024; Watanabe et al., 2013a; Watanabe et al., 2013b), kiss and run (Newton et al., 2006) and bulk endocytosis (Clayton et al., 2009; Kokotos and Cousin, 2015; Kokotos and Low, 2015; Nguyen et al., 2012).

Studies using a variety of model systems, stimulation paradigms and perturbation strategies have produced differing models of dynamin function during SV recycling. Experiments using dominant-negative dynamin mutants, pharmacological disruption of dynamin’s GTPase activity, or peptide blockers of its binding interactions suggest that dynamin is essential for all forms of membrane retrieval (Cheung and Cousin, 2019; Newton et al., 2006; Nguyen et al., 2012). In mammals, three dynamin genes play both distinct and overlapping roles in SV recycling. Combined knockout of dynamin 1 and 3 severely attenuates clathrin-mediated recycling without abolishing it, leading to the proposal that dynamin isoforms act interchangeably and that their collective abundance, rather than any unique function, sets recycling efficiency (Raimondi et al., 2011). Bulk endocytosis, by contrast, continues and is even enhanced in dynamin 1 and 3 double knockouts (Wu et al., 2014), and removal of all three dynamin genes impairs endocytosis during high-frequency stimulation, while leaving substantial recycling intact during sparse or spontaneous activity (Afuwape et al., 2025). RNAi knockdown studies reach different conclusions: depletion of dynamin 1 and 3 slows vesicle protein retrieval under bulk endocytosis conditions (Kononenko et al., 2014), while depletion of individual dynamin genes in sympathetic neurons produces isoform-specific effects on synaptic vesicle recycling tuned to stimulation frequency (Tanifuji et al., 2013). Whether or not these differences reflect genuinely distinct roles for dynamin across cell types and endocytic modes, or instead reflect the different manner in which dynamin has been perturbed, remains unresolved.

All three vertebrate dynamin genes are alternatively spliced, generating an additional potential source of functional diversity (Cao et al., 1998). Of particular interest are seven variants across dynamin 1, 2, and 3 PRDs, which show distinct expression patterns and functional properties. Notably, two dynamin 1 PRD variants (Dyn1xA and Dyn1xB) exhibit different recruitment dynamics to the plasma membrane (Jiang et al., 2024) and function in different types of SV endocytosis: the longer isoform (Dyn1xA) mediates ultrafast endocytosis (Imoto et al., 2022; Imoto et al., 2024) and the shorter isoform (Dyn1xB) facilitates bulk endocytosis and conversion of bulk membranes into SVs (Cheung and Cousin, 2019; Xue et al., 2011). However, because these studies relied on overexpression and peptide inhibitors, and five additional PRDs are largely uncharacterized, the full contribution of distinct dynamin PRDs at mammalian synapses remains unclear.

*Drosophila melanogaster* offers a simplified system for dissecting the roles of dynamin PRD isoforms. The *Drosophila* genome encodes a single dynamin gene, *shibire* (*shi*) that is alternatively spliced to produce long (Shi-L) and short (Shi-S) isoforms differing by a 48 amino acid C-terminal extension in the PRD (Chen et al., 1992; Chen et al., 1991; van der Bliek and Meyerowitz, 1991). As discussed above for mammalian dynamin, there are diverse models for Shi function during SV recycling. Shi was originally thought to be essential for all forms of membrane retrieval based on experiments using temperature sensitive Shi mutants (*shi[ts])* (Delgado et al., 2000; Estes et al., 1996; Kasprowicz et al., 2014; Koenig and Ikeda, 1983, 1989; van de Goor et al., 1995). *shi[ts]* mutants carry a dominant allele, which blocks synaptic transmission at the restrictive temperature (>29°C) by trapping Shi oligomers in aggregate assemblies that acquire biochemical behaviors beyond the wild-type protein (gain of function) while simultaneously sequestering wild-type Shi into trapped complexes (dominant-negative) (Chen et al., 2002; Ikeda et al., 1976; Poodry et al., 1973). Further, *shi[ts]* mutants cause cellular abnormalities at the permissive temperature (Gonzalez-Bellido et al., 2009). These properties raise the possibility that *shi[ts]* phenotypes arise from gain-of-function and/or chronic effects, complicating interpretation of normal Shi function. Studies using photo-inactivated Shi or pan-*shi* RNAi have suggested that Shi is dispensable for (and may even slow) bulk membrane retrieval, and instead regulates SV reformation (Kasprowicz et al., 2014). Overall, conclusions differ both within and between mammalian and *Drosophila* systems, obscuring how dynamin acts during SV recycling. Whether these discrepancies reflect technical limitations (incomplete, indirect, or chronic perturbations) or real biological differences (across species, cell types, and modes of retrieval) remains unresolved.

Mammalian dynamin isoforms with distinct PRDs facilitate different modes of endocytosis, raising the possibility that the two PRDs encoded by the *shi* gene also promote distinct functions in SV recycling. However, despite the long history of studying Shi, the functions of its endogenous isoforms have not been determined. Early studies suggested isoform-specific activities exist, based on differential fractionation from fly head homogenates of 92 and 94 kDa proteins that likely correspond to Shi-S and Shi-L (Gass et al., 1995). Further evidence for isoform-specific functions comes from incomplete rescue of *shi* null mutant lethality and *shi[ts]* paralysis phenotypes by individual isoform cDNAs, with the caveat that the dominant negative *shi[ts]* mutant likely counteracted the rescuing transgene (Staples and Ramaswami, 1999). Here, we use isoform-specific knockdowns and SV recycling assays to investigate the function of Shi-L and Shi-S in the *Drosophila* nervous system.

## Results

### Shi-S and Shi-L PRD isoforms are differentially expressed in the *Drosophila* nervous system

The *shibire* gene locus produces seven unique proteins that can be classified into two groups based on the length of their PRDs. Shi-S consists of three proteins that contain a single unique C-terminal arginine, encoded by a Shi-S specific exon, and Shi-L consists of four proteins that contain a 48 amino acid C-terminal extension encoded by a Shi-L-specific exon (Chen et al., 1992; Chen et al., 1991; van der Bliek and Meyerowitz, 1991) **(Figure 1A-B)**. This 48 amino acid region contains predicted or experimentally validated SH3 domain-binding, actin-binding and phosphorylation sites that may confer unique functions to this class of isoforms **(Figure 1A**, and see “Sequence Analysis” Methods). Notably, neither the Shi-S nor Shi-L isoforms contain an obvious calcineurin-binding motif, which has been shown to specify functions in Dyn1xB (Cheung and Cousin, 2019; Xue et al., 2011).

**Figure 1:**
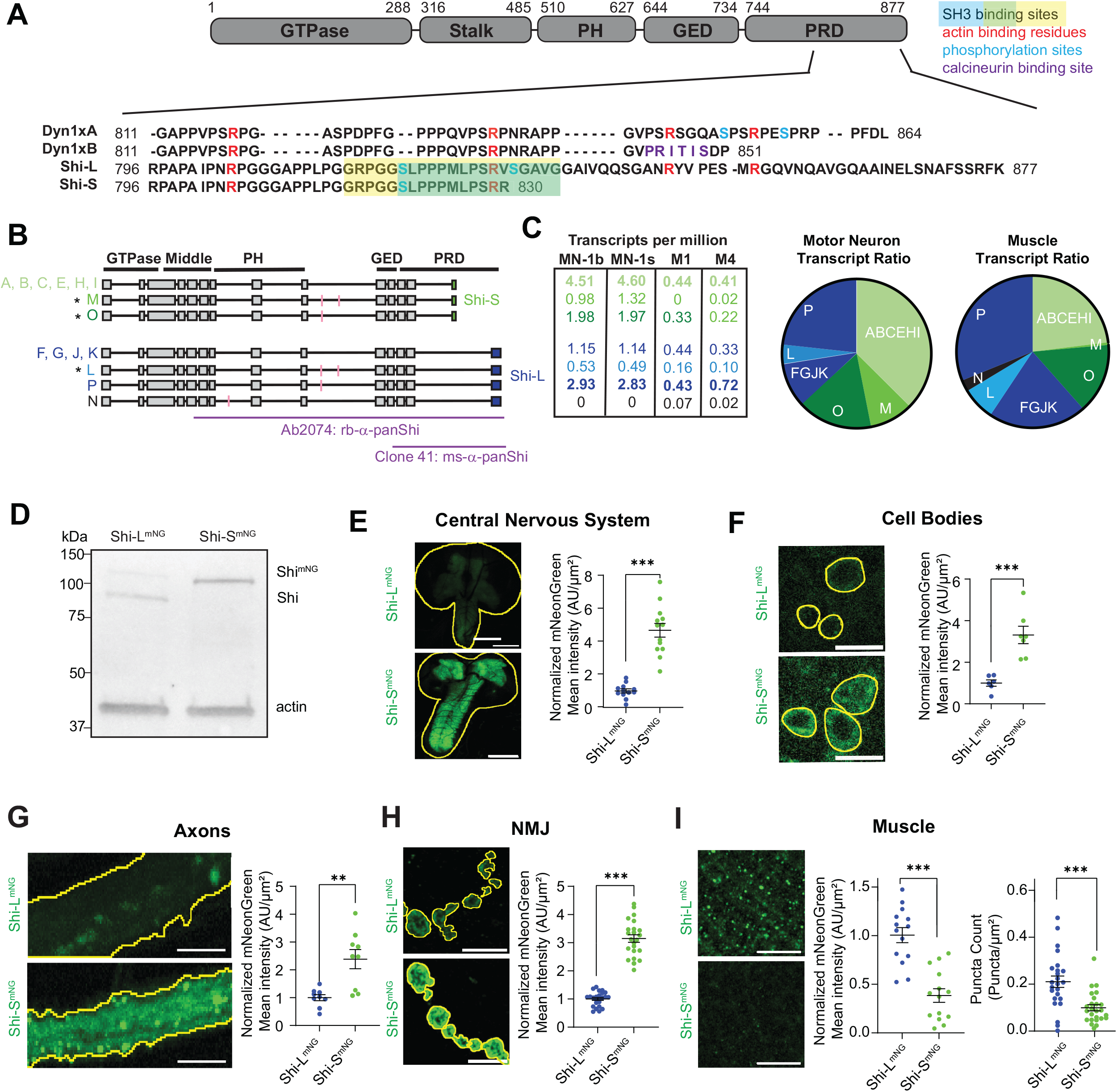
Shi-L and Shi-S are differentially expressed in the *Drosophila* nervous system. **(A)** Sequence alignment of a portion of the PRDs of two isoforms of dynamin 1 (Dyn1xA and Dyn1xB) and two isoforms of Shi (Shi-L and Shi-S). SH3 domain binding sites that partially reside in the C-terminal extension of Shi-L are highlighted. Putative actin binding sites are shown in red. Phosphorylation sites are shown in blue. Calcineurin binding site is shown in purple. See “Sequence Analysis” methods for how these sites were identified. **(B)** Shi splice variants encode 7 unique proteins, with the final exon encoding either Shi-L (blue) or Shi-S (green). Transcripts are further distinguished by their 5’ and 3’ UTRs (not shown). Exons are indicated by grey boxes or magenta lines and introns by horizontal black lines. * marks the isoforms tested in (Staples and Ramaswami, 1999). Purple lines show antigens for dynamin antibodies used in this study. **(C)** Table shows transcripts per million (TPMs) from (Jetti et al., 2023) for each splice variant in MN1-Ib and MN-ISN-1s neurons and M1 and M4 muscles. **(D)** Western blot labeled with rabbit-α-mNG antibody showing levels of Shi^mNG^ in adult *Drosophila* heads. **(E-H)** Maximum intensity projections (MaxIPs) and quantifications of mNG intensity in **(E)** larval central nervous system (optic lobes and ventral nerve cord) (scale bar 100 µm), **(F)** cell bodies (scale bar 10 µm), **(G)** axons (scale bar 5 µm), **(H)** muscle 4 NMJs (Type 1b boutons) (scale bar 5 µm), and **(I)** muscle 6 or 7 (scale bar 5 µm). For **(E-G)**, images were collected on a spinning disk confocal microscope. For **(H-I)**, images were collected on an Airyscan microscope. All graphs show mean +/− SEM. Yellow lines indicate the region of interest for measurements. N = single brain **(E)**, average intensity of cell bodies per larvae **(F)**, average intensity of axons per larvae **(G),** single NMJ **(H)**, single muscle **(I)**. See **Supplemental Table 3** for detailed genotypes, sample sizes, and statistical analyses.

To understand the neuronal functions of Shi PRD isoforms, we started by investigating their relative abundance. We first mined a single-cell RNA-seq dataset of type Ib motor neurons (Jetti et al., 2023), and found six of the seven predicted classes of coding sequences, with Shi-S transcripts (∼63%) more abundant than Shi-L transcripts (∼37%) **(Figure 1C).** By contrast, in larval muscles Shi-L transcripts (∼62%) are more abundant than Shi-S transcripts (∼38%) (Jetti et al., 2023), suggesting that isoform expression is controlled in a tissue-specific manner **(Figure 1C)**. We then used CRISPR and homology-directed repair to generate separate *Drosophila* knockin lines in which the Shi-L and Shi-S-specific exons were seamlessly tagged immediately before their stop codons with mNeonGreen (mNG), preserving their 3’UTRs. Shi-L^mNG^ and Shi-S^mNG^ flies were viable and fertile, and we did not observe paralysis at 38°C **(Supplemental Table 1)**.

Using the mNeonGreen tagged knockin lines, we characterized the levels and distribution of Shi-L^mNG^ and Shi-S^mNG^ in the *Drosophila* nervous system. We found that Shi-S^mNG^ is three to four times more abundant than Shi-L^mNG^ in adult heads, and in larval brain, cell bodies, axons and NMJ **(Figure 1D-H)**. In contrast, Shi-L^mNG^ is more abundant than Shi-S^mNG^ in larval muscle **(Figure 1I).** This result is consistent with their relative transcript levels in these cell types (Jetti et al., 2023). We also found that GAL4/UAS-driven overexpression of Shi-L-eGFP (PJ isoform) or Shi-S-eGFP (PA isoform) in motor neurons led to a modest (1.1-1.4-fold) increase in total Shi immunostaining, and resulted in comparable levels of GFP-tagged protein in larval cell bodies **(Supplemental Figure 1A-B)** and NMJs **(Supplemental Figure 1D-E)**. Moreover, pan-neuronal overexpression yielded similar levels in adult heads by western blot **(Supplemental Figure 1C)**. These results suggest that the endogenous 3:1 ratio of Shi-S: Shi-L in the nervous system is due to regulation of the Shi promoter or UTR rather than regulation at the protein level.

### Shi isoforms have overlapping and distinct distributions at the *Drosophila* NMJ

These observations led us to ask if, in addition to their different expression levels, Shi-L^mNG^ and Shi-S^mNG^ have different subcellular localizations. We recently developed an image analysis pipeline to describe the localization of endocytic proteins to the periactive zone, which forms a mesh approximately surrounding active zones (Del Signore et al., 2023) **(Figure 2A)**. Using Stimulated Emission Depletion (STED) super-resolution microscopy, we found that Shi-L^mNG^ and Shi-S^mNG^ have distinct localization patterns at the periactive zone, as evidenced by a greater proportion of Shi-L^mNG^ compared to Shi-S^mNG^ in the mesh relative to the active zone-containing core (mesh ratio) **(Figure 2B-C).** Further, Shi-L^mNG^ shows a higher coefficient of variation (COV) than Shi-S^mNG^, suggesting that it is more heterogeneously distributed **(Figure 2D)**. However, Shi-L^mNG^ and Shi-S^mNG^ show similar values for a metric of spottiness **(Figure 2E)**. To understand how each isoform localizes relative to the entire population of Shi, we calculated the ratio of isoform-specific signal (using mNG antibodies) to total Shi signal (using pan-Shi antibodies) for each pixel in the presynaptic region. When we plotted how often each ratio value occurred, the values formed a single peak for both Shi-L^mNG^ and Shi-S^mNG^ **(Figure 2F)**. The distribution for Shi-S^mNG^ was shifted toward higher values relative to Shi-L^mNG^, consistent with Shi-S being the more abundant isoform in the nervous system. However, the Shi-L^mNG^ peak was broader than the Shi-S^mNG^ peak, indicating that Shi-L and Shi-S exist in different ratios in different regions of the bouton, and thus may have distinct functions.

**Figure 2:**
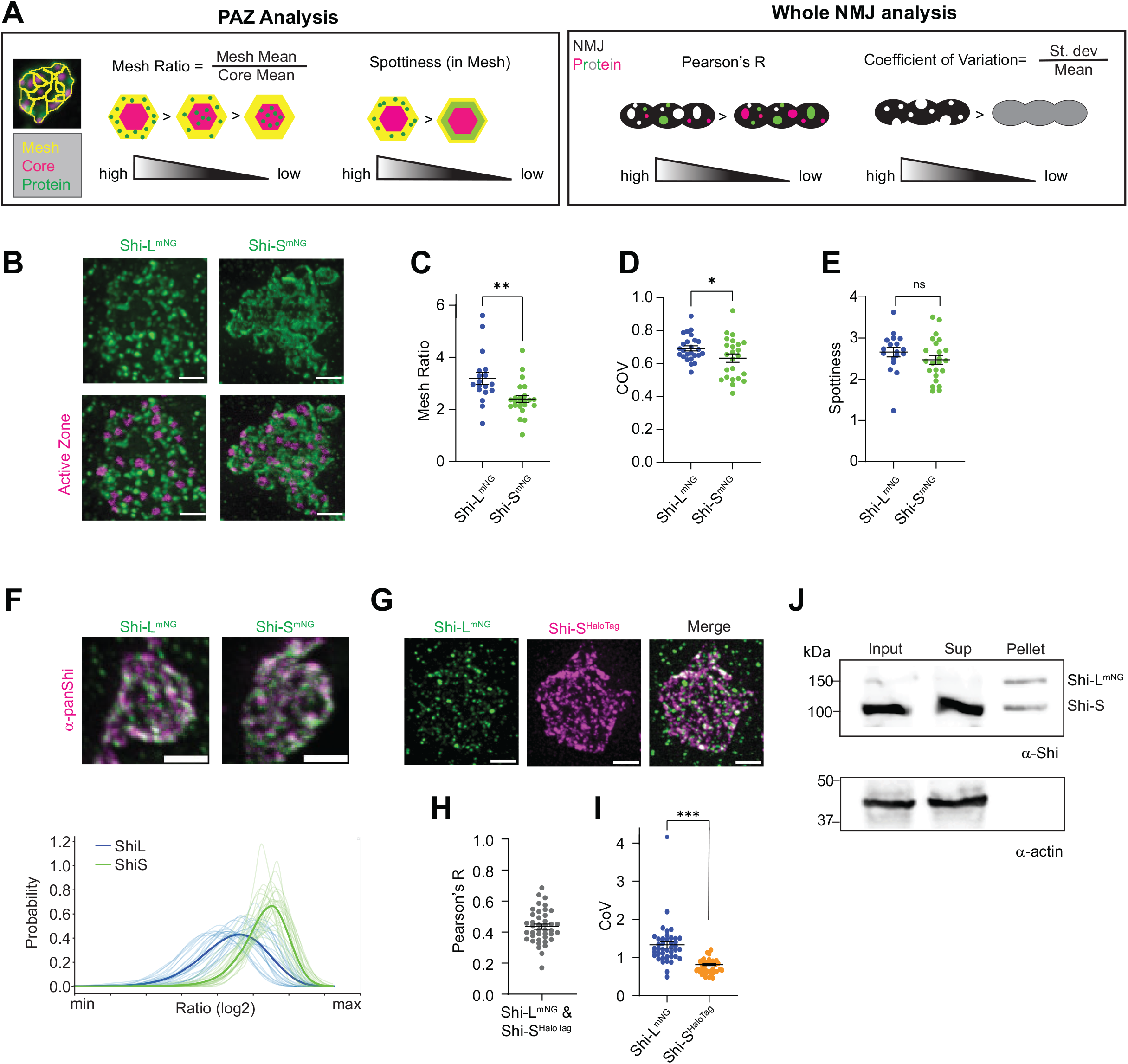
Shi-L and Shi-S have distinct localizations in the *Drosophila* NMJ. **(A)** Cartoon depicting how Mesh Ratio, Spottiness, Pearson’s R (colocalization), and Coefficient of Variation (COV) reflect protein distribution in the presynaptic periactive zone. Mesh Ratio and Spottiness were quantified as previously described (Del Signore et al 2023). Pearson’s R (from 3D stacks) and COV (from 2D projections) were quantified throughout the whole NMJ. **(B)** STED microscopy MaxIPs of muscle 4 NMJs in Shi-L ^mNG^ and Shi-S ^mNG^ larvae stained with chicken-α-mNG and α-BRP to label active zones. (**C-E)** Quantification of protein distribution for the images in **(B**); graphs show mean +/− SEM. **(F)** STED microscopy MaxIPs of muscle 4 NMJs in Shi-L ^mNG^ and Shi-S ^mNG^ larvae stained with chicken-α-mNG and mouse-α-pan Shi to label total Shi protein. Graph shows the ratio of isoform-specific signal to pan-Shi signal, calculated for each pixel in the 3D masked volume and displayed as a probability distribution function (PDF). Bold line indicates average PDF. Since absolute pixel values for α-mNG signal compared to α-pan Shi signal are arbitrary, the ratio is shown on an arbitrary minimum to maximum scale. **(G)** STED microscopy MaxIPs of muscle 4 NMJs in larvae expressing Shi-L ^mNG^ (stained with mouse-α-mNG) and Shi-S^HaloTag^ (stained with HALO dye). (**H-I)** Quantification of protein distribution for the images in **(G)**; graphs show mean +/− SEM. **(B-I)** Shi-L ^mNG^ and Shi-S ^mNG^ intensities were individually adjusted for display. All scale bars are 1 µm. N= single NMJ. **(J)** Shi-L^mNG^ and its binding partners were isolated from fly head homogenates using mNeonGreen beads. The pellet was resuspended in 1/10 original lysate volume of denaturing sample buffer and the same volume of input, supernatant and pellet were loaded on an SDS-PAGE gel. See **Supplemental Table 3** for detailed genotypes, sample sizes, and statistical analyses.

To compare the localizations of Shi-L and Shi-S more directly we used CRISPR and homology-directed repair to generate a *Drosophila* knockin line in which the Shi-S-specific exons were tagged immediately before their stop codons with a HaloTag, preserving their 3’UTRs. In contrast to the Shi-S^mNG^ lines, we were unable to recover Shi-S^HaloTag^ homozygous females or hemizygous males **(Supplemental Table 1)**. Lethality was also observed for flies heterozygous for Shi-S^HaloTag^ and null mutant of *shi*; however, this lethality was rescued by a third chromosome duplication containing the Shi gene (Venken et al., 2010), indicating that lethality of Shi-S^HaloTag^ was not due to a second-site mutation. Thus, while the mNeonGreen tags on Shi are functional, the HaloTag (or different linker used) impairs function. Nevertheless, Shi-S^HaloTag^ was present at the NMJ **(Figure 2G)**. Using STED microscopy in female larvae heterozygous for Shi-L^mNG^ and Shi-S^HaloTag^, we found that Shi-L^mNG^ and Shi-S^HaloTag^ significantly colocalize at the NMJ **(Figure 2H-I)**. However, Shi-L^mNG^ exhibits a punctate distribution whereas Shi-S^HaloTag^ localizes to a smoother periactive zone mesh structure. This is reflected in the difference in the coefficient of variation (COV), where a high COV indicates more heterogeneity **(Figure 2I)**. These differences in subcellular localization suggest that Shi-L and Shi-S could have distinct functions.

Given that Shi-L and Shi-S significantly overlap in their localization patterns at the NMJ **(Figure 2H)**, we considered the possibility that they might form hetero-oligomers and/or co-polymers. Indeed, dynamin oligomerizes via its stalk domain (Antonny et al., 2016) and Shi-L and Shi-S share identical stalks. To test whether Shi-L and Shi-S can co-assemble *in vivo*, we precipitated Shi-L^mNG^ from fly head extracts using α-mNG nanobody-conjugated beads, under conditions that favor higher-order oligomerization (50 mM KCl, (Hinshaw and Schmid, 1995)). Untagged Shi-S co-precipitated with Shi-L^mNG^ in a 1:1 ratio **(Figure 2J)**, suggesting that Shi-L-Shi-S hetero-oligomers are favored over Shi-L-Shi-L homo-oligomers, and that Shi-L and Shi-S may function together. To test direct binding between Shi-L and Shi-S isoforms, we separately purified and labeled Shi-L-SNAP^549^ and Shi-S-SNAP^488^ **(Supplemental Figure 2A-B)** and performed fluorescence colocalization experiments using *in vitro* TIRF microscopy. We pre-incubated Shi-L-SNAP^549^ and Shi-S-SNAP^488^ under conditions that favor the formation of dynamin rings (50 mM KCl, 1 mM MgCl_2_ (Hinshaw and Schmid, 1995)) and imaged the resulting assemblies (**Supplemental Figure 2C)**. Purified Shi-L-SNAP^549^ and Shi-S-SNAP^488^ colocalized in what are likely to be higher-order assemblies (i.e. greater than dimers or tetramers) based on their high fluorescence (∼2000-50,000 AU per channel) and continuous photobleaching (rather than discernable step photobleaching) **(Supplemental Figure 2D-G)**. These results align with previous studies demonstrating co-oligomerization of dynamin isoforms (Altschuler et al., 1998; Barylko et al., 2010; Okamoto et al., 1999; Xie et al., 2012) and further support the idea that Shi-L and Shi-S are capable of functioning together.

### Neuronal Shi-S is essential for viability

To assess the individual contributions of the Shi-L and Shi-S isoforms to synaptic function, we next performed isoform-specific knockdowns using a UAS-driven short hairpin RNAi against mNG. We found that ubiquitous and pan-neuronal knockdown of Shi-S^mNG^ resulted in early lethality, whereas knockdown of Shi-L^mNG^ did not **(Supplemental Table 1)**. As would be expected based on this result, pan neuronal knockdown of both isoforms (with RNAi targeting a common 5’ exon) also resulted in lethality. This suggests that the contribution of neuronal Shi to viability stems primarily from Shi-S function or its higher abundance.

As previously observed for *shi* null mutants (Staples and Ramaswami, 1999), Shi-S^mNG^ knockdown lethality could not be rescued by GAL4/UAS-driven overexpression of one splice variant of Shi-S-eGFP (Shi-PA) or Shi-L-eGFP (Shi-PJ) in the nervous system **(Supplemental Table 2)**. However, we cannot rule out that: (1) another splice variant might rescue lethality or (2) the eGFP tag on the overexpression construct impacted a Shi function required for viability. Interestingly, overexpression of Shi-S-eGFP (Shi-PA) in the nervous system partially rescued lethality due to pan-neuronal knockdown of both isoforms. This may be explained by less efficient knockdown using this pan-Shi RNAi tool compared to mNG-RNAi, especially in the presence of two UAS transgenes. However, given that UAS-Shi-L-eGFP and UAS-Shi-S-eGFP are overexpressed at similar levels **(Supplemental Figure 1)**, this supports the hypothesis that Shi-S-specific functions contribute to viability.

### Depletion of individual Shi isoforms does not cause gross synapse organization defects or compensation by the other isoform

Since the lethality from pan-neuronal knockdown of Shi-S precluded us from exploring its synaptic functions, we next explored the use of more restricted neuronal GAL4 drivers. We found that we could effectively knock down either Shi-L^mNG^ or Shi-S^mNG^ without lethality using Vglut-GAL4, which is expressed in glutamatergic neurons (**Figure 3A, C-D**) (Daniels et al., 2008). Notably, we were also able to achieve pan-Shi knockdown with this driver (albeit less complete than individual isoform mNG knockdowns) to assess the effects of losing both isoforms **(Figure 3A, B)**. NMJs where one or both Shi isoforms were knocked down appeared to be morphologically normal and showed no significant difference in the total number of Type I boutons at muscle 6/7 NMJs compared to control (mCherry RNAi) animals **(Supplemental Figure 3A).** Moreover, these NMJs did not exhibit any defects in the distribution of active zones labeled by Bruchpilot (BRP) **(Supplemental Figure 3B-E**). It has been proposed that dynamin is an organizer of the periactive zone due to its ability to interact with many ligands via its PRD (Kasprowicz et al., 2014; Raimondi et al., 2011). However, after knockdown of either Shi-L or Shi-S we did not detect any dramatic defects in the levels or organization of the periactive zone proteins Nervous Wreck (Nwk-mammalian homolog is FCHSD2) and Dynamin associated protein 160kDa (Dap160-mammalian homolog is intersectin) **(Supplemental Figure 4)**.

**Figure 3:**
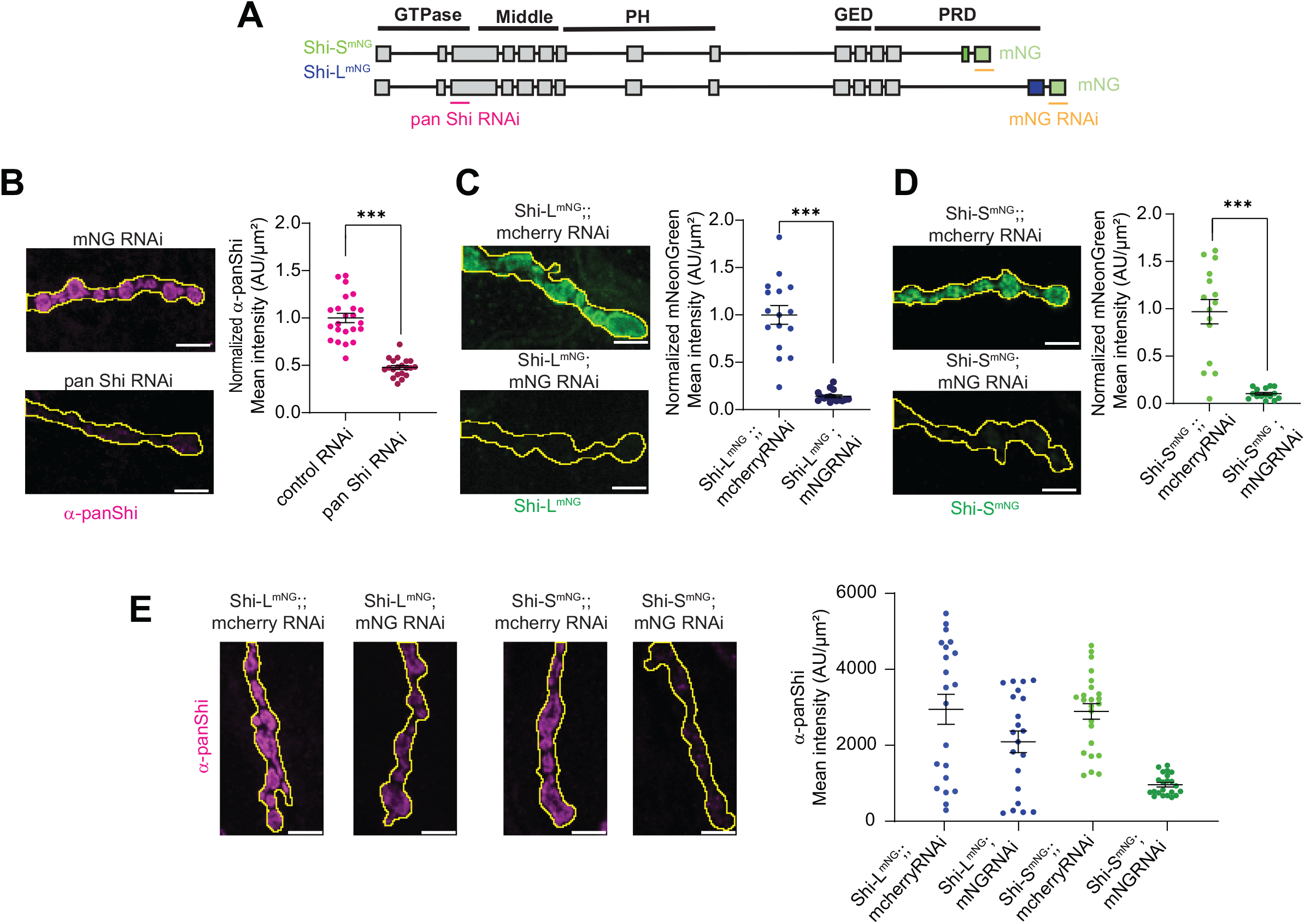
Efficient knockdown of Shi-L^mNG^ and Shi-S^mNG^ at the NMJ. UAS-mNG RNAi or control mcherry RNAi were expressed under the control of the motor neuron driver GAL4^Vglut^. **(A)** Schematic showing Shi exons targeted by pan Shi RNAi and mNG RNAi. Spinning disk MaxIPs of muscle 4 NMJs in controls and **(B)** pan Shi knockdowns stained with rabbit-α-pan Shi, **(C)** Shi-L^mNG^ knockdowns and **(D)** Shi-S^mNG^ knockdowns. **(E)** Spinning disk MaxIPs of muscle 4 NMJs in controls and isoform knockdowns stained with mouse-α-pan Shi. Note there is no compensation relative to expected knockdown levels. All scale bars are 5 µm, and graphs show mean intensity +/− SEM. N=single NMJ. See **Supplemental Table 3** for detailed genotypes, sample sizes, and statistical analyses.

To determine whether depletion of one isoform might have triggered compensatory changes in the levels of the other, we measured total Shi protein levels at the NMJs of Shi-L^mNG^ or Shi-S^mNG^ knockdown animals using a pan-Shi antibody **(Figure 3A, E)**. Knockdown of Shi-L^mNG^ led to a 29% reduction in total Shi levels, while knockdown of Shi-S^mNG^ resulted in an 81% reduction (**Figure 3E**). These findings align with the relative abundance of the isoforms in the nervous system **(Figure 1D-H)** and suggest that there is little or no compensation upon depletion of either isoform.

### Shi-S is required for high-demand endocytosis

Using the isoform-specific knockdowns, we next investigated how Shi isoforms function during SV recycling. First, we asked whether synapses in which Shi had been knocked down could sustain high-frequency neurotransmitter release, by measuring the excitatory junction potential (EJP) when the neuron was stimulated at 10 Hz for 5 min in 2 mM CaCl_2_. This stimulation protocol elicits extensive SV exocytosis and requires efficient SV recycling, likely by a clathrin-mediated mechanism, to maintain the SV pool (Heerssen et al., 2008; Kasprowicz et al., 2008; Kasprowicz et al., 2014; Verstreken et al., 2008). Shi knockdowns exhibited distinct phenotypes: pan Shi RNAi (loss of both Shi-L and Shi-S) and Shi-L^mNG^ knockdown NMJs displayed only a mild reduction in EJP amplitude over time compared to the initial response, and this decline was not significantly different compared to controls or between genotypes **(Figure 4A)**. However, knockdown of Shi-S^mNG^ resulted in strong synaptic depression, as shown by the dramatic decline in EJP amplitude **(Figure 4A).** This aligns with what was observed in *shi[ts]* mutants at the restrictive temperature (Delgado et al., 2000; Ikeda et al., 1976; Koenig and Ikeda, 1983) and upon photoinactivation of Shi (Kasprowicz et al., 2014). These results suggest that loss of Shi-S leads to defects in synaptic vesicle reformation. After a one-minute rest and stimulation at 0.1 Hz for 200 seconds, all genotypes showed robust recovery from high frequency stimulation, as shown by the EJP amplitude returning to baseline **(Figure 4A).** This efficient recovery in Shi-S^mNG^ knockdowns suggests that under low frequency stimulation, a new readily releasable pool of vesicles can be generated by the remaining Shi-L, or through a Shi-independent mechanism.

**Figure 4:**
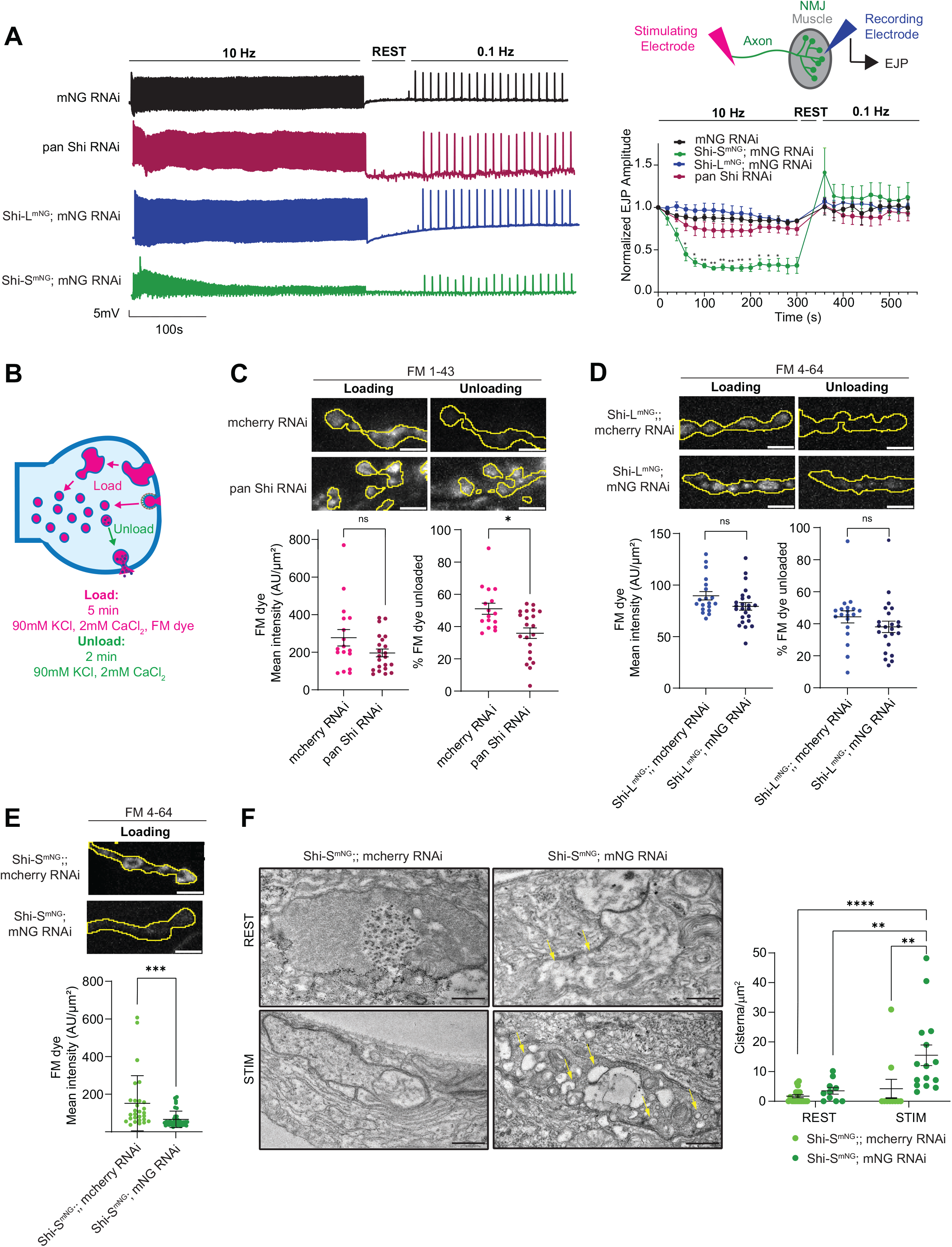
Shi-S^mNG^ knockdown disrupts SV endocytosis in response to high frequency stimulation. **(A)** Representative electrophysiological recordings from NMJs of controls and pan Shi, Shi-L^mNG^, and Shi-S^mNG^ knockdowns. The experimental setup for electrophysiology experiments is shown in the top right. A stimulating electrode suctions an axon to administer stimuli and a recording electrode inserted into the postsynaptic muscle monitors excitatory junction potential (EJP). NMJs in 2 mM CaCl_2_ were stimulated at 10Hz for 5 min followed by a 1-minute rest and stimulation at 0.1Hz for 200 seconds. Graph shows EJP amplitude (mean +/− SEM) over time normalized to the EJP amplitude at timepoint 0 for each genotype. * indicates significance relative to the same timepoint in controls (black). **(B)** Schematic of FM dye uptake assays. Spinning disk MaxIPs of muscle 6/7 NMJs in controls and **(C)** pan Shi knockdowns (raised at 29°C) **(D)** Shi-L^mNG^ knockdowns (raised at 25° C) and **(E)** Shi-S^mNG^ knockdowns (raised at 25°C) in FM uptake experiments. NMJs were stimulated with 90 mM KCl and 2 mM CaCl_2_ for 5 minutes to load with FM dye (magenta), imaged, stimulated again with 90 mM KCl and 2 mM CaCl_2_ for 2 minutes to unload the dye and then imaged a second time. All scale bars are 5µm. Graphs show mean uptake (FM dye mean intensity +/− SEM) and release (percent FM dye unloaded +/− SEM). **(F)** Transmission electron microscopy Images of muscle 6/7 boutons from controls and Shi-S^mNG^ knockdowns at rest or stimulated with 90 mM KCl and 2 mM CaCl_2_ for 10 minutes. Scale bar 400 nm. Graph shows the number of bulk cisternae (>80nm in diameter) per unit area (mean +/− SEM). N single NMJ **(A-E)** and single bouton **(F)**. See **Supplemental Table 3** for detailed genotypes, sample sizes, and statistical analyses.

During periods of high exocytic demand, bulk endocytosis retrieves membrane by internalizing large plasma-membrane cisternae from which synaptic vesicles are subsequently reformed (Kokotos and Cousin, 2015). To investigate the roles of Shi-L and Shi-S isoforms in this process, we conducted FM dye uptake assays using a 90 mM KCl stimulation paradigm to evoke synaptic vesicle release and bulk endocytic uptake at the NMJ **(Figure 4B)**. We found that pan Shi knockdowns did not exhibit a significant defect in dye loading but were unable to unload the dye following a second stimulus **(Figure 4C)**, as previously observed (Kasprowicz et al., 2014). Since internalized FM dye was observed in aberrant intracellular structures in the above-mentioned study, we expect that this unloading defect stems from a failure to resolve synaptic vesicles from bulk structures. Shi-L^mNG^ knockdown NMJs loaded and unloaded a similar amount of dye to controls **(Figure 4D)**. In contrast, loss of Shi-S^mNG^ resulted in a severe loading defect **(Figure 4E)**, similar to phenotypes observed in *shi[ts]* mutants at the restrictive temperature (Kuromi and Kidokoro, 1998). We were unable to conduct further unloading experiments in Shi-S^mNG^ knockdown animals due to the loading defect.

We considered two possible biological explanations for lack of loading: Either bulk endocytosis fails to initiate, or bulk invaginations form but remain on the membrane due to failed scission. To distinguish between these two models, we performed transmission electron microscopy on synapses stimulated with high KCl solution. Similar to what was previously observed using photoinactivated Shi mutants (Kasprowicz et al., 2014), we found that Shi-S^mNG^ knockdown led to an increase in the number of bulk cisterna (>80 nm in diameter) per unit area in boutons stimulated with 90mM K+ for 10 minutes **(Figure 4F).** Given that FM dye could not be loaded into boutons of the same genotype experiencing a similar stimulation paradigm **(Figure 4E)**, we expect that these bulk cisternae are still attached to the plasma membrane and therefore any dye in these structures was removed in the wash step.

The dramatic phenotypes in Shi-S^mNG^ knockdowns and lack thereof in Shi-L^mNG^ and pan Shi knockdowns suggest several interpretations. One possibility is that our pan Shi RNAi is not sufficient to evoke a dramatic phenotype because it only partially depletes Shi-S (as suggested by our immunostaining experiments, **Figure 3B**). Alternatively, the Shi-L that remains in the Shi-S^mNG^ knockdowns could act in an inhibitory manner to block bulk uptake. The latter would align with the prediction that Shi can slow bulk uptake (Kasprowicz et al., 2014). If Shi-L had an inhibitory function, we expect that overexpression of Shi-L-eGFP would block FM dye uptake; however, we observed no difference in dye loading or unloading upon overexpression of Shi-L-eGFP in motor neurons **(Supplemental Figure 5A-B).** This result supports the model that mild phenotypes in pan Shi RNAi conditions stem from inefficient knockdown.

### Shi-S abundance drives its function during SV recycling

Together, the above assays show that Shi-S (but not Shi-L) is necessary and sufficient to facilitate SV recycling during intense stimulation. This may be due to specific properties of the Shi-S PRD or due to its higher abundance compared to Shi-L. To distinguish between these two models, we performed rescue experiments by overexpressing Shi-L-eGFP or Shi-S-eGFP in motor neurons in which Shi-S^mNG^ was knocked down. First, we confirmed efficient knockdown and overexpression in these larvae **(Supplemental Figure 6A-B)**. We noted that mNeonGreen RNAi does not impact the Shi-eGFP transgenes and confirmed that these transgenes are only expressed at 1.1-1.4 fold compared to endogenous Shi levels **(Supplemental Figure 1)**. Then, we measured the EJP amplitude in response to 10 Hz stimulation for 5 min in 2 mM CaCl_2_. Upon high frequency stimulation, the dramatic decrease in EJP amplitude caused by Shi-S^mNG^ knockdown could be rescued by expression of either Shi-L-eGFP or Shi-S-eGFP **(Figure 5A)**, suggesting that the abundance of Shi, as opposed to specific contributions from its unique PRD, is the primary contributor to its functions in SV recycling.

**Figure 5:**
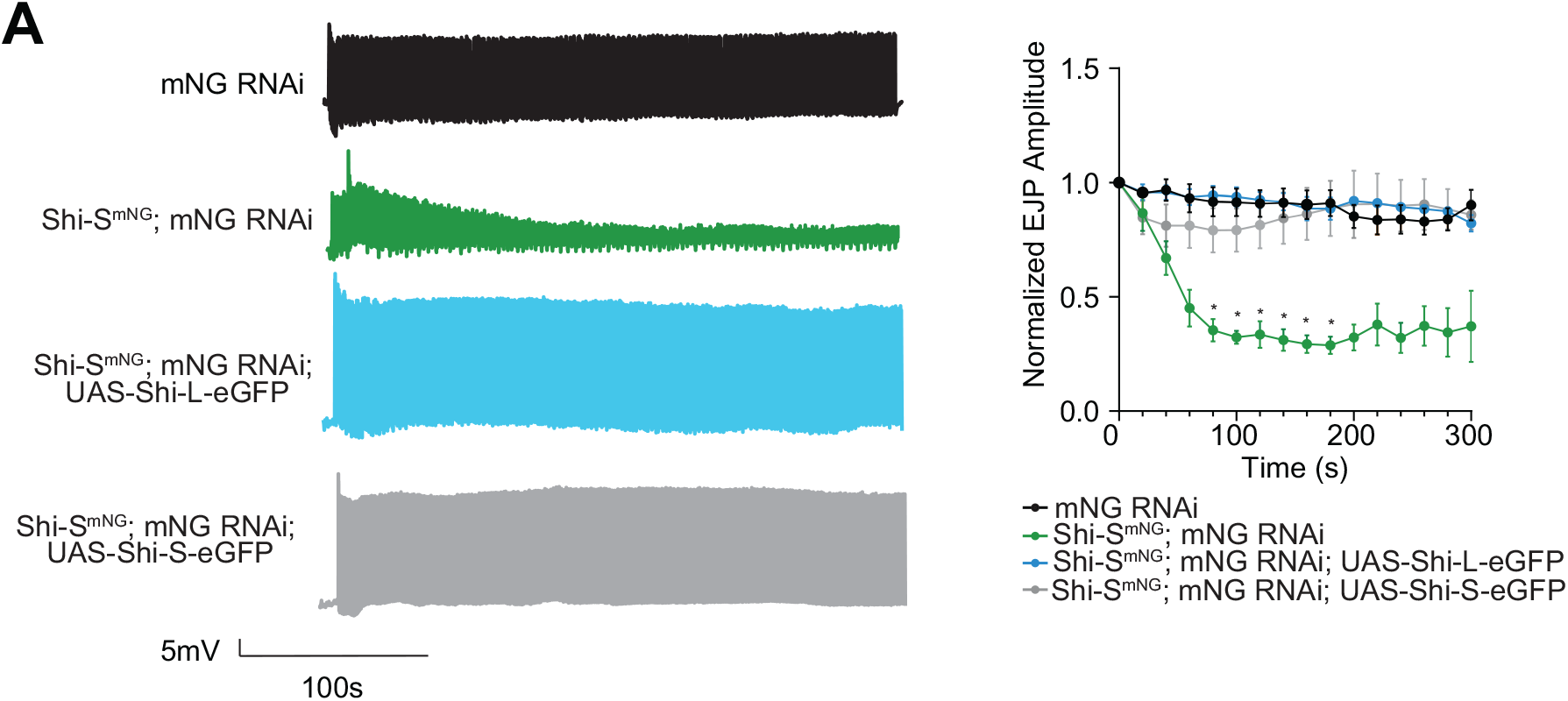
Shi-S abundance is the main contributor to its functions in SV recycling. **(A)** Traces of electrophysiological recordings at NMJs in controls and Shi-S^mNG^ knockdowns overexpressing either Shi-L-eGFP or Shi-S-eGFP. NMJs in 2 mM CaCl_2_ were stimulated at 10Hz for 5 min. Graph shows the EJP amplitude (mean +/− SEM) over time normalized to the EJP amplitude at timepoint 0 for each genotype. The control and Shi-S^mNG^ knockdowns are the same as Figure 4A but were reanalyzed with less stringent detection parameters that provided better detection for the rescue conditions. * indicates significance relative to the same timepoint in controls (black). See **Supplemental Table 3** for detailed genotypes, sample sizes, and statistical analyses.

### Shi-S regulates SV recycling during spontaneous activity and low intensity stimulation

Next, we tested whether Shi-S played a role in regulating spontaneous activity or activity in response to low intensity stimulation (as opposed to high intensity stimulation in the experiments above). We first compared miniature EJP (mEJP) amplitude and frequency at the NMJs of Shi-S^mNG^ control and knockdown animals in 0.4 mM CaCl_2_ (**Figure 6A**). At the Shi-S knockdown NMJs, there was an increase in mEJP amplitude **(Figure 6B)** and mEJP frequency **(Figure 6C)**, suggesting that the amount of neurotransmitter per SV increases, and that SVs spontaneously fuse with the plasma membrane more often compared to control NMJs. We then stimulated the same NMJs at 1 Hz for 200 seconds in 0.4 mM CaCl_2_ and found that EJP amplitude decreased in Shi-S^mNG^ knockdowns **(Figure 6D)**. Using the mEJP and EJP amplitudes, we also calculated the quantal content, or the number of vesicles released per action potential (Bykhovskaia and Vasin, 2017). Similar to *shi[ts]* mutants (Narayanan et al., 2005), the quantal content decreased in Shi-S^mNG^ knockdowns **(Figure 6E)**, indicating that fewer vesicles were released in response to sparse action potentials. Thus, Shi-S is necessary for multiple aspects of synaptic vesicle cycling and release, even under mild stimulation conditions or at rest.

**Figure 6:**
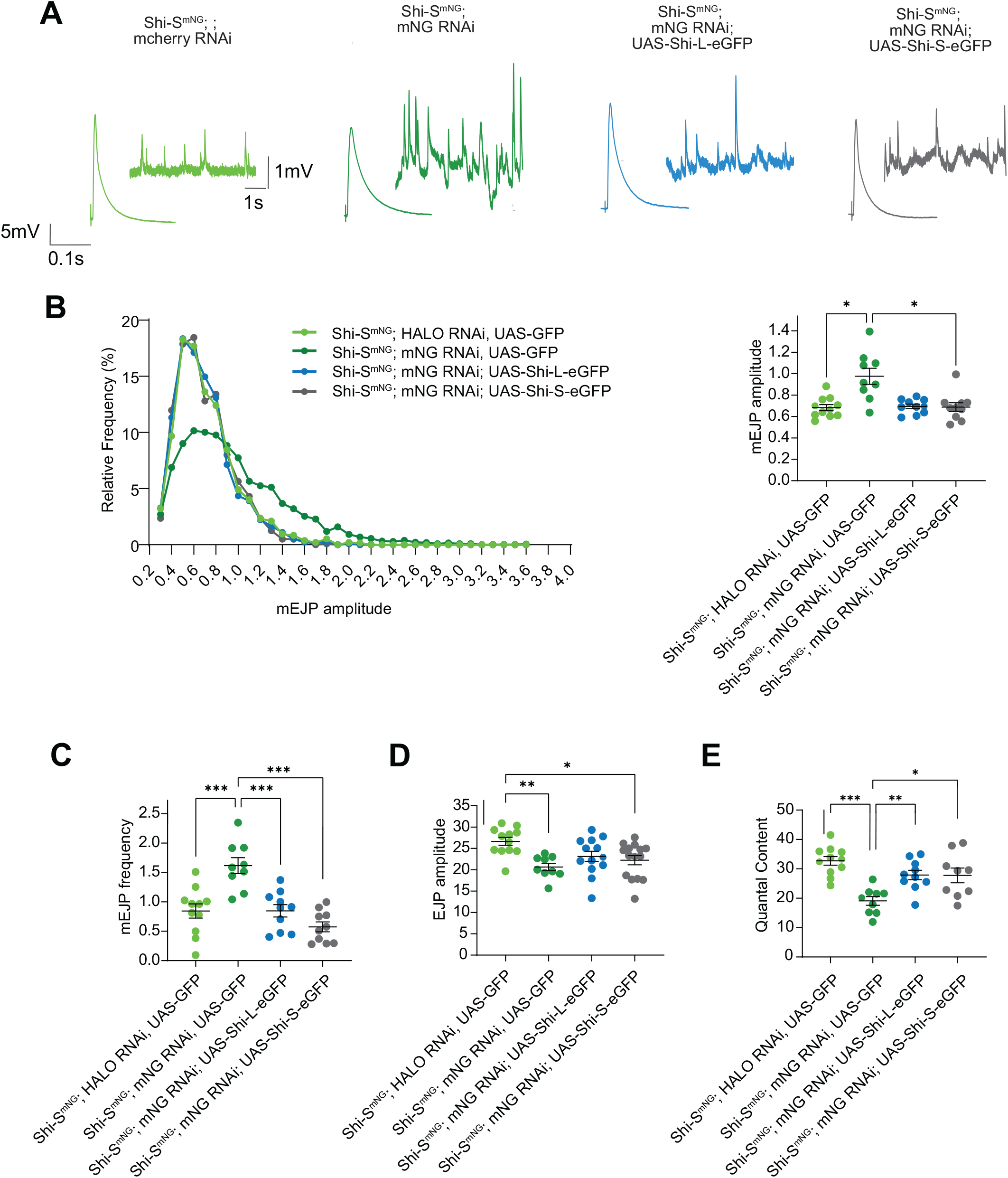
Shi-S abundance is the main contributor to its functions in SV recycling during spontaneous activity and under low intensity stimulation. **(A)** mEJP trace set in EJP trace for controls and Shi-S^mNG^ knockdowns when GFP, Shi-L-eGFP and Shi-S-eGFP are overexpressed. **(B-E)** Recordings were performed in 0.4 mM CaCl_2_ on muscle 6/7 NMJs in controls and Shi-S^mNG^ knockdowns with GFP, Shi-L-eGFP and Shi-S-eGFP overexpression. EJP amplitude was recorded at 1 Hz. Graphs show **(B)** mEJP amplitude, **(C)** mEJP frequency, **(D)** EJP amplitude and **(E)** quantal content (mean +/− SEM). N=single NMJ. In histogram in B, N= single mEJP. See **Supplemental Table 3** for detailed genotypes, sample sizes, and statistical analyses.

Finally, we asked whether Shi-S^mNG^ function during spontaneous activity and low intensity stimulation was due to specific properties of the PRD or its higher abundance compared to Shi-L. To answer this question, we again performed rescue experiments by overexpressing Shi-L-eGFP or Shi-S-eGFP in motor neurons where Shi-S^mNG^ was knocked down. Expression of either Shi-L-eGFP or Shi-S-eGFP at least partially rescued the increase in mEJP amplitude **(Figure 6B)** and mEJP frequency **(Figure 6C),** and the decrease in EJP amplitude **(Figure 6D)** and quantal content **(Figure 6E)** caused by Shi-S^mNG^ knockdown. These results suggest that the abundance of Shi, rather than its specific PRD, is the main contributor to its functions in SV recycling at rest and in response to low intensity stimulation, as was the case in response to high intensity stimulation.

## Discussion

Here we show that long and short C-terminus isoforms of *Drosophila* dynamin/Shibire make unequal contributions to synaptic vesicle recycling at the larval neuromuscular junction. By endogenously tagging each isoform, we found that Shi-S is expressed at substantially higher levels than Shi-L, and that the two isoforms have overlapping and distinct patterns of localization at the periactive zone. Depletion of Shi-S (but not Shi-L) disrupted bulk endocytic uptake and the reformation of synaptic vesicles under high-intensity stimulation. At rest and under low-frequency stimulation, Shi-S knockdown phenotypes were more nuanced: spontaneous release was elevated, and evoked responses were reduced. Surprisingly, overexpression of either the Shi-L or Shi-S isoform could rescue the defects caused by Shi-S depletion, suggesting that abundance, rather than PRD identity, is the principal determinant of the Shi isoform roles. This result is unexpected in light of mammalian studies, in which distinct PRD variants have been proposed to direct dynamin into specific modes of endocytosis through isoform-specific interactions (Cheung and Cousin, 2019; Imoto et al., 2022; Imoto et al., 2024; Jiang et al., 2024; Xue et al., 2011).

Our findings instead suggest that the critical roles of dynamin during synaptic vesicle recycling do not require a specialized isoform, or differentially interacting PRD, but simply a sufficient quantity of either Shi isoform. Why neurons express both isoforms, and at a particular ratio, rather than simply expressing a high level of Shi-S or Shi-L may be for other reasons (discussed below).

### Shi abundance shapes multiple modes of synaptic vesicle recycling

Our approach of knocking down Shi-S, which removed ∼80% of total Shi and left ∼20% intact (largely as untargeted Shi-L), produced a set of informative phenotypes that help clarify long-standing debates about which forms of membrane retrieval require dynamin. Unlike *shi[ts]*, our knockdown removes protein rather than poisoning it, avoiding the confounds of dominant-negative tools. It achieves greater depletion than the tools used previously in *Drosophila*: pan-Shi RNAi reached only ∼50% knockdown both here and in earlier work, and acute photoinactivation phenocopied this RNAi suggesting partial inhibition (Kasprowicz et al., 2014). Our depletion also occupies a useful middle ground in timing: because the regulatory elements we used to express mNeonGreen RNAi activate late in embryonic development (Mahr and Aberle, 2006; Sprecher et al., 2004), Shi is likely progressively lost over the ∼4 days of larval development. This is more acute than germline knockout in mice or weeks-long depletion in culture (Afuwape et al., 2025; Raimondi et al., 2011), and we show that this does not result in compensatory upregulation of the untargeted isoform. This window biases our perturbations toward functions requiring ongoing maintenance of Shi levels, such as SV recycling, while processes established earlier in development, when residual protein was more abundant, may be spared. Therefore, our Shi-S knockdown represents a severe regime where enough dynamin is lost to identify which steps of the synaptic vesicle cycle are most sensitive to its abundance, without impacting its functions in development.

We found that under high-frequency stimulation (10 Hz) in Shi-S knockdowns, the amplitude of evoked release decreased rapidly over time. This observation is consistent with results across other experimental systems showing that sustained high-frequency transmission depends on abundant dynamin to regenerate release-competent vesicles (Afuwape et al., 2025; Delgado et al., 2000; Ferguson et al., 2007; Ikeda et al., 1976; Kasprowicz et al., 2014; Koenig and Ikeda, 1983; Kononenko et al., 2014; Raimondi et al., 2011). In addition, we found that release recovered rapidly once the train ended, indicating that vesicles can still reform once demand falls. This result is consistent with previous studies reporting little or no endocytic defect in dynamin mutants under low-demand conditions, and could reflect retrieval via residual Shi-L or a dynamin-independent route (Afuwape et al., 2025; Ferguson et al., 2007; Kononenko et al., 2014; Raimondi et al., 2011).

At low-frequency stimulation and at rest, Shi-S knockdown led to additional informative phenotypes. The persistence of spontaneous events and the rapid recovery of EJPs in the knockdowns (described above) indicate that a substantial fraction of vesicle recycling proceeds independently of Shi-S, and of abundant Shi more generally. Notably, spontaneous release was not merely preserved: events were both larger in amplitude and more frequent. Because our knockdown is cell-autonomous to the presynaptic neuron, the amplitude increase most likely reflects a presynaptic change such as increased vesicle size, consistent with the larger or more variable quantal size reported in mammalian dynamin mutants (Ferguson et al., 2007; Raimondi et al., 2011); unfortunately vesicles were too densely packed in our EM to accurately measure their size.

The frequency increase, by contrast, is different from the reduced or unchanged spontaneous rate seen after dynamin loss in mammals (Afuwape et al., 2025; Raimondi et al., 2011), as well as in other *Drosophila* mutants like photoinactivated Shi (Kasprowicz et al., 2014) and *shi[ts]* (Koenig and Ikeda, 1983). Our results point to a functional difference in these vesicles or their release machinery upon loss of Shi-S. This could occur through a direct effect of Shi on vesicle sorting and composition or an indirect, homeostatic effect on the release apparatus. Given evidence that spontaneous and evoked retrieval are partially spatially and functionally segregated at the larval NMJ (Sabeva et al., 2017), one possibility is that vesicles reformed under low-Shi conditions are biased toward spontaneous fusion. This raises the interesting hypothesis that cells could use Shi regulation to control spontaneous release.

In contrast to the increase in mEJP amplitude and frequency, we found that evoked release and quantal content were reduced in Shi-S knockdowns, similar to dynamin 1 and 3 knockouts (Afuwape et al., 2025; Ferguson et al., 2007; Raimondi et al., 2011). That evoked release is impaired while spontaneous release is enhanced points to a specific reduction in the evoked-competent pool, which may be caused by fewer readily-releasable vesicles (Afuwape et al., 2025), altered vesicle composition (Raimondi et al., 2011), or both. Notably, we did not address the role of Shi in ultrafast endocytosis or kiss and run retrieval, modes for which there is little evidence at this synapse and which our assays are not designed to detect (Dickman et al., 2005; Gan and Watanabe, 2018; Heerssen et al., 2008; Kasprowicz et al., 2008; Verstreken et al., 2002).

Finally, our finding that Shi-S knockdown disrupts bulk endocytosis helps to clarify a long-standing question in the field: whether dynamin is required for bulk membrane uptake or only for the reformation of vesicles from bulk endosomes. Under high potassium stimulation Shi-S knockdown NMJs still internalized membrane (since bulk cisternae were readily apparent by EM), but fail to retain FM dye after washing, suggesting that cisternae remain continuous with the plasma membrane. These results cannot be explained by an inhibitory role for the residual Shi-L, since Shi-L overexpression does not impair FM uptake. Our findings are different from Wu et al. (Wu et al., 2014), where bulk cisternae in Dyn1 or Dyn1/3 knockouts retained a fluid-phase marker and were sealed off from the plasma membrane, raising the possibility that residual Dyn2 was sufficient to pinch cisternae from the surface. This framework also reconciles apparently conflicting results between our findings with previous work in *Drosophila*.

Kasprowicz et al. (Kasprowicz et al., 2014) found that synapses with ∼50% Shi RNAi or acute Shi photoinactivation retained FM dye in wash-resistant cisternae that were refractory to subsequent unloading and concluded that Shi is dispensable for bulk uptake but required for vesicle reformation. We suspect this reflects the partial depletion of Shi in these studies: enough dynamin remained (50% by RNAi, and by its phenocopy a comparable fraction after photoinactivation) to pinch cisternae off the membrane, whereas the ∼20% Shi-L left in our knockdown cannot. Together these results suggest an abundance-dependent hierarchy of dynamin function in bulk endocytosis. As dynamin is progressively depleted, reformation of synaptic vesicles from cisternae is the first function to fail at ∼50% depletion; pinching of cisternae from the plasma membrane is more resistant to partial loss of dynamin, failing only under the more severe depletion that leaves ∼20% Shi-L; and initial membrane uptake is the most resistant, persisting even with only Shi-L. Importantly, neither membrane uptake nor pinching required a direct calcineurin-binding motif of the kind implicated in mammalian bulk endocytosis (Cheung and Cousin, 2019; Xue et al., 2011). By contrast to these loss-of-function manipulations, *shi[ts]* exerts a different effect: it blocks even initial membrane uptake, likely due to gain-of-function effects at the internalization step.

### Isoform abundance, not PRD identity, determines Shi’s role in synaptic vesicle recycling

The three mammalian dynamin genes differ along their entire length, and have distinct GTPase activities (Warnock et al., 1997), curvature sensing and generation capabilities (Liu et al., 2011), membrane interactions, expression levels (Cao et al., 1998; Sontag et al., 1994), and localizations (Cao et al., 1998; Gray et al., 2003; Liu et al., 2008; Okamoto et al., 2001; Raimondi et al., 2011). Any functional differences among them, such as their frequency-tuned roles in vesicle replenishment (Tanifuji et al., 2013) or their non-redundant contributions at synapses (Ferguson et al., 2007) or non-neuronal cells (Bhave et al., 2020) could arise from any of these properties and need not reflect a specialization of the PRD. Splice variants of a single gene provide a more direct test of PRD specialization, because they share every other domain and differ only in the C-terminal tail that carries isoform-specific interactions.

Specialized roles for the mammalian variants (i.e. Dyn1xA in ultrafast endocytosis, Dyn1xB in bulk endocytosis) have been established through overexpression rescue of a knockout, peptide inhibition, or imaging of tagged constructs (Cheung and Cousin, 2019; Imoto et al., 2022; Imoto et al., 2024; Jiang et al., 2024; Xue et al., 2011), and each was tested largely against a single endocytic mode in cells in culture. By contrast, our endogenously tagged knockins and knockdowns allow us to evaluate levels and localization of one isoform at its native level and deplete it while leaving the other intact, in the context of an intact organism.

We found that Shi-L and Shi-S are both expressed in the nervous system, with Shi-S as the more abundant isoform, and that this ratio is reversed in muscle. Given that the neuronal ratio was lost when overexpressing Shi isoforms from a heterologous UAS promoter, it is likely that this ratio reflects tissue-specific regulation of the Shi promoter or untranslated regions rather than control at the protein level. The isoforms also exhibit partial co-localization at the NMJ, form a complex with one another in head extracts, and can associate *in vitro*, consistent with the co-oligomerization of dynamins 1, 2, and 3 in mammals (Altschuler et al., 1998; Barylko et al., 2010; Okamoto et al., 1999). We found that their localizations are nonetheless partially distinct (as reported for mammalian isoforms (Cao et al., 1998)), likely due to isoform-specific interactions through the 48–amino acid Shi-L extension. Our finding that the isoforms are both expressed in neurons and occupy distinct regions suggests that the Shi-L extension can support isoform-specific functions.

We assayed the consequences of isoform depletion across the range of SV recycling modes and found that Shi isoforms did not exhibit specialized functions, since either supports recycling when supplied at sufficient levels in the Shi-S knockdown background. Shi-S therefore predominates in SV recycling not because of its short PRD, but because it is the more abundant isoform. That dynamin abundance governs its contribution to recycling has been shown for whole genes (Afuwape et al., 2025; Raimondi et al., 2011). Our results extend this conclusion to splice isoforms and indicate that the variables encoded in this region (PRD ligand binding, actin binding, phosphorylation) do not specify an activity critical for SV recycling. Because the isoforms can assemble into mixed oligomers, this suggests that a functional assembly requires only enough total dynamin and is largely independent of the ratio of tails within it.

Importantly, these results do not imply that the PRD tail is inert. In mammals, tail-specific interactions route dynamin to particular modes of endocytosis, including endophilin binding for ultrafast endocytosis, and calcineurin-binding for bulk endocytosis (Cheung and Cousin, 2019; Imoto et al., 2024; Xue et al., 2011). What we can conclude is that a dedicated long-tail or calcineurin-binding–dependent pathway is not a central component of the SV cycle in our experimental system.

### Shi abundance and isoform-specificity are not essential for presynaptic organization or morphogenesis

Our recent work shows that endocytic proteins are targeted to the periactive zone largely independently of active-zone machinery and activity (Emperador-Melero et al., 2026), leaving open the question of how they are recruited and organized near synapses. Dynamin makes many scaffolding or regulatory interactions with SH3 domain-containing endocytic proteins (Meinecke et al., 2013; Rosendale et al., 2019; Sundborger and Hinshaw, 2014) and has been proposed to influence the abundance and organization of endocytic proteins in neurons (Kasprowicz et al., 2014; Raimondi et al., 2011), making it good candidate for organizing the periactive zone. Endocytic proteins have also been implicated in synaptic development by controlling the traffic of signaling receptors that instruct expansion of the NMJ onto the muscle as it grows during larval development, as well as the size and spacing of active zones (Dickman et al., 2006; Goel et al., 2019; O’Connor-Giles et al., 2008; Rodal et al., 2008). Our results indicate that neither Shi abundance nor a specific isoform is required for recruitment or organization of periactive zone components Nwk and Dap160 or for synaptic growth or active zone size or distribution.

This indicates either (1) that dynamin is not required for these processes; (2) that ∼20% residual protein may suffice for organization even where it is insufficient for SV recycling or (3) that synaptic architecture may be established earlier in development, when perduring Shi is more abundant (which is intriguing as it would indicate that periactive zone architecture is stable over many hours or days).

### Conclusions

If the Shi isoforms are interchangeable for SV recycling, why are two maintained? In mammals, dynamin isoform-specific functions reside in the tail, and the Shi-L extension likewise carries predicted actin-binding, SH3-binding, and phosphorylation sites that could recruit distinct partners or be differentially modified. The conservation of PRD isoforms across arthropods argues that they are important: diverse insects such as mosquito (*Anopheles gambiae*) and beetle (*Tribolium castaneum*), and chelicerates such as the spider *Trichonephila clavata*, all encode two similar PRDs. These properties point to roles we did not assay. For example, Shi functions in the dense-core-vesicle pathway (Wong et al., 2015) (where isoform contributions are untested). Shi-L also regulates the actin cytoskeleton during myoblast fusion (Zhang et al., 2020) in muscles where Shi-L predominates and dynamin is known to be critical, as underscored by DNM2-linked human centronuclear myopathies (Zhao et al., 2018). Further, while either isoform restored recycling in the Shi-S knockdown, neither cDNA transgene restored viability, suggesting a requirement beyond the recycling functions we assayed. This lack of rescue could also reflect properties outside the tail that are required beyond SV recycling, since our transgenes each contained only one of three small variable insertions between the PH and GED domains that could affect membrane binding or catalysis. Overall, our findings support the conclusion that how much dynamin is present, more than which isoform, governs most of its function in the synaptic vesicle cycle. The fact that distinct PRD isoforms are conserved across hundreds of millions of years of evolution implies that the alternative tails serve functions we have yet to define, most likely beyond the synaptic vesicle cycle itself.

## Methods

### Drosophila culture

Flies were cultured using standard media and techniques. All flies were maintained at low density and raised at 25°C or 29°C (RNAi experiments were raised at 29 degrees unless otherwise noted). Wandering third instar larvae were used in all experiments unless otherwise noted. Either male or female larvae were used for experiments, with each independent experiment using only one sex. See **Supplemental Table 3** for detailed sex and genotype information for each experiment. Unless otherwise noted, all experiments used Vglut-Gal4 as the driver.

### Drosophila strains

*Drosophila* strains used in this study can be found in **Supplemental Table 4**. Strains made in this study were created as follows:

#### Knockins

An mNeonGreen tag or HALO7 tag was knocked into the endogenous *shi* locus by CRISPR-mediated gene editing (performed by WellGenetics Inc. using modified methods of (Kondo and Ueda, 2013)). A flexible linker was added between Shi-S final exon and the mNeonGreen tag. In brief, a gRNA sequence for Shi-L (TCAGTTCGCGATTCAAGTAA[TGG]) or Shi-S (ACATCTACATATGTATCTAG[GGG]) was cloned into a U6 promoter plasmid. This targets the stop codon in exon 14 and labels all Shi-L isoforms (F, G, J, K, L, P, N) or all Shi-S isoforms (A, B, C, E, H, I, M, O). Two homology arms and either cassette mNeonGreen 3xP3 RFP, which contains mNeonGreen and a floxed 3xP3 RFP, or cassette Halo7 3xP3 RFP, which contains Halo7 and a floxed 3xP3 RFP, were cloned into pUC57 Kan as donor template for repair. shi/CG18102 targeting gRNAs and heat shock Cas9 were supplied in DNA plasmids. These constructs were microinjected into embryos of the control strain w[1118]. CRISPR generates a break in shi/CG18102 and is replaced by cassette mNeonGreen 3xP3-RFP or Halo7 3xP3 RFP. F1 flies carrying 3xP3 RFP selection marker were further validated by genomic PCR and sequencing. Finally, the floxed 3xP3-RFP cassette was excised to generate the knockin strain used in experimental crosses.

#### RNAi

mNeonGreen RNAi and HaloTag RNAi lines were created in accordance with the Drosophila TRiP project (Perkins et al., 2015). In brief, a foldback RNA was designed in the MiR1 scaffold and cloned between the NheI and EcoR1 sites in pUASzMiR (for germline and soma expression, 10X UAS) as described (Ni et al., 2011). For mNeonGreen, the top strand oligo was ctagcagtAGACCGAGCTCAACTTCAAGGtagttatattcaagcataCCTTGAAGTTGAGCTCGGTCTgcg and the bottom strand oligo was aattcgcAGACCGAGCTCAACTTCAAGGtatgcttgaatataactaCCTTGAAGTTG AGCTCGGTCTactg. For HaloTag, the top strand oligo was ctagcagtAGCTGATCATCGATCAG AACGtagttatattcaagcataCGTTCTGATCGATGATCAGCTgcg and the bottom strand oligo was aattcgcAGCTGATCATCGATCAGAACGtatgcttgaatataactaCGTTCTGATCGATGATCAGCTactg. pUASzMiR-mNeonGreen or pUASzMir-HaloTag were inserted into the attp40 locus (Markstein et al., 2008) on *Drosophila* chromosome II by BestGene Inc.

#### UAS-Shi-EGFP

Shi-L (PJ) or Shi-S (PA) were cloned by Azenta into pBI UASc-eGFP (Wang et al., 2012). Plasmids were injected into y1 w67c23; P{CaryP}attP2 (FBst0008622) (by BestGene Inc). Transformants were identified by the w+ marker and balanced.

### Immunohistochemistry

Wandering third instar larvae were dissected in calcium-free HL3.1 saline (Feng et al., 2004) in Sylgard dishes. Larval fillets were fixed in HL3.1 containing 4% formaldehyde for 10-20 minutes. In high K+ stimulation experiments, wandering third instar larvae were dissected in calcium-free HL3 in Sylgard dishes that were never exposed to fixative. Axons were cut proximal to the ventral nerve cord and then larvae were shifted to high K+ HL3 containing 90 mM KCl and 2 mM CaCl_2_ for 10 minutes. Control animals experienced mock stimulation in standard HL3 for 10 minutes. Larvae underwent three quick (1 second) washes with HL3 and then fixed in HL3 containing 4% formaldehyde for 15 minutes.

Fixed samples were rinsed three times in washing buffer (PBS+0.1% Triton X100) followed by three ten-minute washes in washing buffer. In experiments using HALO dye, larvae were incubated in 500 nM HALO-JF549 (Promega, Inc.) in washing buffer for 15 minutes followed by another round of three ten-minute washes in washing buffer. Fixed and permeabilized larvae were incubated in a blocking solution containing 3% BSA (weight/volume) in 0.1% Triton X100 in PBS for 30-60 min. Then larvae were incubated with primary antibody in blocking solution either overnight at 4°C or 4 hours at room temperature. Primary antibody details can be found in **Supplemental Table 5**. Samples were washed in washing buffer and incubated for two hours at room temperature with dye-conjugated secondary antibodies or α-HRP antibodies in the appropriate washing buffer. The secondary antibodies were conjugated to Alexa 488 (used at 1:250), Rhodamine Red-X (used at 1:500), Alexa 647 (1:250) (Jackson Immunoresearch), STAR red (used at 1:200) or STAR orange (1:200) (Abberior). Larvae were washed then mounted in Prolong Diamond mounting medium, Vectashield or Abberior Liquid Mount Antifade.

### Image acquisition

Segments A2-A4 of muscle 6/7 or muscle 4 were imaged, as specified in figure legends. All images were acquired at room temperature, using spinning disk confocal, Airyscan or STED microscopes, as specified in figure legends and detailed below. Images for each independent experiment were acquired using the same settings for all conditions.

#### Spinning disk confocal

Images were collected using Nikon Elements AR software on a Nikon Ni-E upright microscope equipped with 10x objective (n.a. 0.3), 60x oil immersion objective (n.a. 1.4), 60x water immersion objective (n.a. 1.0), a Yokogawa CSU-W1 spinning disk head, and an Andor iXon 897U EMCCD camera. Images of brains were collected using the 10x objective and 1 μm step size. Images of fixed cell bodies, axons and NMJs were collected using the 60x objective and 0.3 μm step size. Images of live NMJs in FM dye experiments were collected on 60x water objective and 0.3 μm step size.

#### Airyscan

Images were acquired using Zen Black acquisition software on a Zeiss880 Fast Airyscan microscope in super resolution acquisition mode. A 63x oil immersion objective (n.a. 1.4) and 0.18 μm step size were used. All raw image stacks were processed in Zen Blue to construct Airyscan images using 3D Airyscan processing with automatic settings.

#### STED

Images were acquired using iMspector Lightbox Software on an Abberior FACILITY line microscope with 60x (n.a. 1.3) silicone immersion objective or 60x (n.a. 1.42) oil immersion objective, pulsed excitation lasers (561 nm and 640 nm), and a pulsed depletion laser (775 nm) to deplete all signals. Images were acquired in 3D STED mode (∼25%) and pixel size was set to 40 nm. The silicon objective was used on samples mounted in Abberior Liquid Mount Antifade and the oil objective was used on samples mounted in Prolong Diamond mounting medium.

### Huygens deconvolution

Images collected on a STED microscope were deconvolved with Huygens Essential Deconvolution software v24.04 (Scientific Volume Imaging, The Netherlands, http://svi.nl). For each experiment, we used a Classic Maximum Likelihood Estimation (CMLE) deconvolution algorithm and applied consistent parameters for each channel imaged. We used standard settings in the Huygens software to automatically determine the theoretical point spread function (PSF) and the optimal number of iterations. We manually measured the background signal and calculated a signal to noise ratio based on measured signal and background intensities for each protein. In some cases, the acuity value was optimized to smooth or sharpen the deconvolved image. Deconvolution was performed on a workstation with an Intel(R) Xeon(R) W-3335 CPU @3.40GHz, 256GB RAM, and an NVIDIA RTX 4000 Ada Generation 20GB GDDR6 GPU.

### Fixed image analysis

#### Bouton counting

Images of muscle 6/7 NMJs were blinded. Type 1 boutons in segment A3 were counted manually. Satellite boutons were defined in accordance with (Dickman et al., 2006b) as extensions of five or fewer boutons emanating from the main branch of the nerve terminal.

#### Intensity, COV and colocalization measurements

##### Cell bodies

A single z-slice containing a group of 2-4 cell bodies belonging to the motor neuron layer of the ventral ganglion (beneath neuropil) was cropped in a 100-pixel x 100-pixel square. Individual cell bodies were outlined manually in FIJI. Mean intensity was measured inside the region of interest. For all images, background was subtracted using the rolling ball method with a radius of 50 pixels.

##### Axon/whole brain

Sum projections were created and portions of the whole brain and individual axons proximal to the ventral ganglion (within 100-300 μm) were outlined manually in FIJI. Mean intensity was measured inside the region of interest. For all images, background was subtracted using the rolling ball method with a radius of 50 pixels.

##### NMJ

To prepare images for analyses, we manually cropped out regions that were obscured by axon bundles. Masks of the NMJ were generated using a presynaptically enriched label (e.g. α-HRP). For mask generation, images were subjected to a Gaussian blur filter and auto thresholded using a thresholding algorithm; Blur radius and the specific threshold algorithms used were empirically optimized for each experiment to consistently and accurately reflect the presynaptic area in different groups (the same settings were used for all NMJs within any given experiment). Signal intensities were measured in 3D using a FIJI script, COV was measured in 2D on maximum intensity projections using a FIJI script, and colocalization analysis was performed in 3D using the Coloc2 plugin for ImageJ or by calculating the Pearson Correlation Coefficient between two arrays. For all images, background was subtracted using the rolling ball method with a radius of 50 pixels.

##### Muscle

To prepare images for analysis, we manually cropped out regions that contained axons, NMJs or non-muscle containing background. Signal intensities were measured in 3D in the entire cropped area using a FIJI script. For all images, background was subtracted using the rolling ball method with a radius of 50 pixels.

##### Muscle spot detection

To prepare images for analysis, we manually cropped out regions that contained NMJ. Then we optimized maxima detection by qualitatively assigning a prominence value at which all obvious spot structures were detected. The prominence value was normalized to the mean intensity in the muscle to calculate the prominence factor (prominence factor= prominence/Mean intensity) for individual muscles. The average prominence factor was calculated for at least 5 muscles and that value was used as the prominence factor in the analysis. In the analysis, images were subjected to a Gaussian blur filter and Mexican Hat LoG Filter. Maxima were detected using the calculated prominence factor. Maxima served as a centroid for the Seeded Region Growing(Adams and Bischof, 1994; Bischof, 1994) which isolates individual spots. Spots above 250 pixels were excluded. The number of spots were counted and normalized to area of the muscle.

#### PAZ Analysis

##### PAZ processing and segmentation

All image processing and data extraction were performed using custom FIJI and Python scripts (Del Signore et al., 2023) and available at https://github.com/rodallab/paz-analysis. First, maximum intensity projections were made of the upper half of NMJ terminals, to analyze a single plasma membrane surface. Second, a 2D mask of the total presynaptic area was generated by summing all channels in each image and then thresholded by intensity using a thresholding algorithm that was determined to accurately detect NMJ shape. Third, a periactive zone mesh composite image was defined by the composite of two PAZ proteins and BRP (subtracting BRP from the sum of PAZ protein signals) or simply PAZ proteins if no BRP was present. Fourth, periactive zone units were detected as local intensity minima and then expanded by the seeded region growing algorithm (via the IJ-Plugins Toolkit). The thresholds for minima detection were computed automatically and periactive zones detected at the edge of boutons (defined as having a mean Euclidean distance map score of less than 7.5 pixels) were excluded from analysis as these are not planar. Settings were applied identically within an experiment. This process segmented the NMJ into ‘mesh’ and ‘core’ regions, where the periactive zone mesh is a ∼175 nm wide band localized on average ∼330 nm from the center, and the ‘core’ region is interior to this periactive zone mesh

##### Quantification of PAZ architecture

PAZ segmentation is used to measure

#### Mesh Ratio

The ratio of the mean mesh intensity over the mean core intensity describes a protein’s localization within a PAZ unit. Mesh ratios greater than 1 indicates the protein is more localized to the mesh. Mesh ratio less than 1 indicates the protein is more localized to the core.

#### Spottiness

Images were processed with a commonly used spot detection filter (Laplacian of Gaussian) and magnitude of response to this filter was calculated as the root-mean-squared value of the filtered image (because the filter ranges from positive to negative across spot boundaries). Higher scores indicate a more punctate or spotty distribution of signal.

#### Colocalization

Pearson’s correlations coefficients (PCC) were measured on 2D half-maximum-intensity projection images. *Active zone Segmentation:* Local maxima were used as seeds to generate objects by the seeded region growing ImageJ plugin (ImageJ-Plugins). Merged active zones were split by a distance transform watershed (Legland et al., 2016).

#### Isoform Ratio

To segment NMJs, a 3D mask was created from the combined isoform-specific and pan-Shi signals, using the Li algorithm in FIJI. The log(2) of the ratio of isoform-specific signal to pan-Shi signal was calculated for each pixel in the 3D masked volume. To analyze the distribution of values, the probability distribution function of pixel ratio values was calculated for each image using scipy.stats gaussian kernel density estimate. Plots show the average probability distribution function (bold) overlaid on the individual traces.

##### Quantification of AZ architecture

Active zone (AZ) segmentation was used to measure:

#### BRP count/NMJ area

The number of BRP puncta segmented as active zone objects was counted and normalized to the area of the NMJ.

#### BRP area/count

The total area of BRP objects was measured and divided by the total number of BRP objects to calculate the average size of an active zone labelled with BRP in each NMJ.

#### BRP mean intensity

The mean intensity of BRP in whole boutons was measured, as described above.

### Electrophysiology

Third-instar larvae were dissected in HL3.1 saline with either 0.4 (Figure 6) or 2 mM CaCl₂ (Figures 4 and 5). Sharp-electrode recordings of neuromuscular junctions were obtained from muscle 6 in abdominal segment A2, A3 or A4 using glass microelectrodes with a resistance of 12–35 MΩ. Stimulation was delivered with an A-M Systems Model 2100 stimulator. Miniature excitatory junctional potentials (mEJPs) and evoked EJPs were recorded using an Axon Instruments MultiClamp 700A amplifier.

Recordings were included for analysis only if the resting membrane potential was more hyperpolarized than −55 mV, the baseline drift was ≤10% and the input resistance exceeded 4 MΩ. Events were detected using custom MATLAB scripts based on code from https://github.com/marderlab/Spike_analysis.

For spontaneous transmission, 0.4 mM CaCl₂ was used and mEJPs were recorded for at least 2 minutes. For low frequency stimulation, 0.4 mM CaCl₂ was used and at least 15 nerve evoked potentials (EJPs) were recorded at 1Hz. To do this, motor axons innervating muscle 6 in the segment were recruited and the average suprathreshold evoked EJP amplitude was empirically identified for each synapse. Quantal content was calculated by dividing the mean EJP by the mean mEJP after correction of EJP amplitude for nonlinear summation, according to previously described methods (Martin, 1955). A reversal potential of 0 mV was used for this correction.

For high-frequency stimulation, 2 mM CaCl₂ was used. The suprathreshold evoked EJP amplitude was empirically identified for each synapse and animals were then stimulated at 0.1 Hz for at least 15 action potentials followed by 10 Hz for 5 minutes. After stimulation, a 1-minute rest was given and recovery of the action potential was tested.

### FM dye labeling

Wandering third instar larvae were dissected in calcium-free HL3 saline, taking care not to overstretch the larval filet. Axons were cut proximal to the ventral nerve cord prior to stimulation. NMJs were stimulated for 5 minutes with an HL3 solution containing 90 mM KCl, 2 mM CaCl_2_ and 4uM of either FM4-64 (Shi-S^mNG^ and Shi-L^mNG^ knockdowns) or FM1-43 (pan Shi knockdown). Larvae underwent three quick (1 second) washes with HL3 saline, followed by a 3-minute wash with 1:1000 advicept in HL3 then three quick (1 second) washes with HL3. Then muscle 6/7 NMJs were imaged on a spinning disk confocal microscope. For Shi-L^mNG^ and pan Shi knockdowns, NMJs were stimulated again with HL3 containing 90 mM KCl and 2 mM CaCl_2_ for 2 minutes to unload the FM dye. Larvae underwent three quick (1 second) washes with HL3 saline. The same muscle 6/7 NMJs from the loading step were imaged. Mean intensities of FM dye in the NMJ following loading and unloading were measured as described above. Loading was quantified as the mean intensity of FM dye following the first stimulation. Unloading was quantified by percent of FM dye unloaded using the formula (mean intensity of loaded dye-mean intensity of dye following unloading)/mean intensity of loaded dye).

### Transmission electron microscopy

Wandering 3rd instar larvae were dissected in HL3 saline in groups of equal numbers of control and mutant animals. Fillets were subjected to stimulation with a high potassium HL3 solution containing 90 mM KCl, 1.5 mM CaCl_2_ for 10 mins, with control animals experiencing mock stimulation in standard HL3 for 10 minutes. Samples were fixed overnight at 4°C in 4% paraformaldehyde, 1% glutaraldehyde in 0.1M sodium cacodylate buffer, pH 7.2. Samples were washed three times for 10 min in 0.1M cacodylate and were then postfixed in 1% Osmium Tetroxide, 1.5% Potassium ferrocyanide in 0.1M cacodylate for 3 hours. Samples were washed for 10 min with 0.1M cacodylate then twice for 10 min with H2O. Then the samples were incubated with 2% aqueous Uranyl Acetate in water for 30 mins. Stepwise dehydration was conducted for 10 minutes each in 30%, 50%, 70%, 85%, 85% and three times in 100% ultra-dry ethanol. Samples were incubated for 1 hour each in 100% Propylenoxide, 3:1 Propylenoxide to 812 TAAB Epon Resin (TAAB Laboratories Equipment Ltd, Aldermaston, Enland), 1:1 Propylenoxide to epon, 1:3 Propylenoxide to epon and left overnight in 100% epon while shaking. The next day, samples were embedded in molds containing new epon and left to polymerize at 60°C for 48 hours. Molds were trimmed with a razor blade prior to sectioning. 70nm thin sections were cut on a Leica UC6 Ultramicrotome (Leica Microsystems, Buffalo Grove, IL), collected on to 2×1mm copper grids coated with formvar and carbon (Electron Microscopy Sciences, Hatfield, PA). Some samples were stained in lead citrate. Grids were imaged using a FEI Morgagni transmission electron microscope (FEI, Hillsboro, OR) operating at 80kV and equipped with a Nanosprint5 CMOS camera (AMT, Woburn, MA) at 4400-5600x magnification.

Analysis of bulk endosomes in electron micrographs was performed in FIJI on blinded images. Only one image per bouton was analyzed, and images were included in analysis if they had sufficient resolution to discern bulk endosomes, a clear bouton boundary, and were greater than 100 nm^2^. An object was considered a bulk endosome if it had a distinct boundary, was not electron-dense, and had a diameter of greater than 80nm, as measured with the line tool in FIJI. The bouton area was measured by manually tracing the bouton boundary. The number of bulk endosomes was then normalized to the bouton area.

### SDS PAGE gels

Purified proteins or heads (5-15 pooled per genotype) from *Drosophila* adults aged 3-10 days were homogenized in 2x Laemmli buffer (10uL per head). Lysate samples were boiled for 1 min and centrifuged. Samples were loaded into BioRad Any KD precast gels and fractionated by gel electrophoresis.

### Western blot

Samples were fractionated by SDS/PAGE and transferred to nitrocellulose membranes. Membranes were blocked with a 4:1 solution of TBST: western blot blocking buffer for 30 minutes-1 hour. Membranes were incubated in primary antibody either overnight at 4°C or room temperature for one hour. This was followed by washing in TBS+0.1% Tween-20 and incubation with dye-conjugated secondary antibodies in TBS+0.1% Tween-20 for 1.5 hours. The secondary antibodies were conjugated to Dylight 488 (used at 1:100), Dylight 680 (used at 1:200), or Dylight800 (1:2000) (Rockland). The blot was subsequently imaged on BioRad Chemidoc system and quantified using FIJI.

### Protein expression and purification

pFastBac1-Shi-L-SNAP or pFastBac1-Shi-S-SNAP **(Supplemental Table 6)** were transformed into DH10EmBacY cells (Trowitzsch et al., 2010) to create a bacmid containing the gene of interest through T7 integration. The bacmid was prepared using the PureLink HiPure Plasmid Midiprep Kit. Presence of the gene of interest was confirmed via PCR using pUC/M13 primers. 1ug of the bacmid was transfected into adherent Sf9 cells cultured in Gibco Sf-900 III SFM media using 10uL of ExpiFectamine™ Sf Transfection Reagent in 250 μL Opti-MEM™ I Reduced Serum Medium. Once signs of viral infection occurred (72-120 hours post transfection) the media containing virus was collected, filtered and stored at 4°C in the dark as P0 viral stock. ∼1mL of P0 virus was added to 50mL of Sf9 cells in suspension, and the same procedure was followed to amplify a P1 viral stock. Following optimization for viral titer, 100uL-1.5mL of P1 viral stock was added to 600mL of Sf9 cells in suspension. Cells were diluted to 1×10^6^ cells/mL every 24 hours for 72 hours. Cells were collected via centrifugation, resuspended in 10mL of cold PBS, snap frozen in liquid nitrogen and stored at −80°C.

For purification, cells were lysed using a sonicator in 2x lysis buffer (2x PBS and 60 mM imidazole pH8, 1x Halt protease inhibitor). Shi constructs were purified via affinity chromatography on an AKTA FPLC using a 1 mL HisTrap column equilibrated in wash buffer (1x PBS and 30 mM imidazole pH8). Protein was eluted using an elution buffer (1x PBS and 500 mM imidazole pH8) and peak fractions were pooled. Eluate was applied to a 10/300 Superdex 200 column for gel filtration into 20 mM HEPES-KOH pH7.3, 150 mM KCl, 1 mM EGTA, 1 mM DTT. Peak fractions were pooled. For SNAP labeling, a 3x molar ratio of SNAP dye (final concentration of 20-30 µM) was added to the pooled fractions and incubated for 24 hours at 4°C. Excess dye was removed using PD-10 desalting columns equilibrated in 20 mM HEPES-KOH pH7.3, 150 mM KCl, 1 mM EGTA, 1 mM DTT. The protein was concentrated, then aliquoted, frozen in liquid nitrogen and stored at −80°C.

### TIRF microscopy

TIRF microscopy images were collected on a Nikon-Ti2000 inverted microscope equipped with a 150-mW argon laser (Melles Griot), a 60× TIRF objective with a NA of 1.49 (Nikon Instruments), and an electron-multiplying charge-coupled device (EMCCD) camera (Andor Ixon, Belfast, Northern Ireland). Focus was maintained by the Perfect Focus system (Nikon Instruments) and settings were applied using imaging software Elements (Nikon Instruments). Images were analyzed in FIJI (National Institutes of Health, Bethesda).

Glass coverslips were cleaned in 2% Hellmanex III solution while sonicating at 50°C for 60 minutes and then rinsed with MilliQ water, 180 proof EtOH and 0.1M KOH. Coverslips were dried with compressed N2 gas and then coated overnight with 40mg/mL PEG-saline solution +/− 8m/mL BioPEG in 200 proof EtOH and 10uL Ultra-pure HCl. Coverslips were then rinsed with MilliQ water, dried with compressed N2 and stored at −80°C. A coated coverslip was fastened to a 30 μL flow cell using double sided tape (Grace Bio-Labs SecureSeal Adhesive Sheets. 17.8cm x 25.4cm x 0.12mm) and epoxy resin to create a TIRF chamber. The coverslip was washed with 100 μL HBSA for 1 minute. HBSA was washed out with 1x TIRF buffer (10 mM imidazole, pH 7.4, 50 mM KCl, 1 mM MgCl_2_, 1 mM ethylene glycol-bis(β-aminoethyl ether)-N,N,N′,N′-tetraacetic acid (EGTA), 0.2 mM ATP, 10 mM DTT, 15 mM glucose, 20 μg/ml catalase, 100 μg/ml glucose oxidase).

#### TIRF imaging

Purified Shi-L-SNAP^549^ and Shi-S-SNAP^488^ in 150 mM KCl, 0 mM MgCl_2_ were mixed in low salt (50 mM KCl, 1 mM MgCl_2_) 1x TIRF buffer for 10 minutes at room temperature. Mixtures were flowed into a bio-PEG TIRF chamber, allowed to passively absorb to the surface, and visualized by TIRF microscopy. The percentage of spots that formed hetero-oligomers and homo-oligomers was manually counted in three independent experiments.

#### Step Photobleaching

Purified Shi-L-SNAP^549^ and Shi-S-SNAP^488^ in 150 mM KCl, 0 mM MgCl_2_ were mixed in low salt (50 mM KCl, 1 mM MgCl_2_) 1x TIRF buffer for 30 minutes at room temperature.

Mixtures were added to a PEG TIRF chamber, allowed to passively absorb to the surface, and visualized by TIRF microscopy. One field of view was continuously photobleached for 5 minutes using maximum laser power. A 5-pixel x 5-pixel square was drawn around hetero-oligomers (where Shi-L and Shi-S were present). The mean intensity of Shi-L and Shi-S in the first frame in each ROI was measured and plotted in a histogram. The mean intensity of each ROI was plotted over time to create a photobleaching trace.

### Pulldowns

Lysates were made from snap-frozen fly heads in lysis buffer (20 mM HEPES, pH 7.5, 50 mM KCl, 0.5 mM EGTA, 1 mM MgCl_2_, and 0.2% (v/v) Igepal CA630) supplemented with Halt protease inhibitor, using a Teflon homogenizer. Lysates were centrifuged at max speed and the non-lipid containing layer was collected. A Bradford assay was performed to calculate the total amount of protein in w1118, Shi-L^mNG^ and Shi-S^mNG^ lysates. Concentrations of the lysates were normalized to the least concentrated lysate. Concentrations of proteins on beads were normalized using empty beads. The bead volume was restricted to two-thirds of the total reaction volume. Normalized lysates were applied to mNeongreen agarose beads for 1 hour at 4°C or SNAP Capture Magnetic Beads for 3 hours at 4°C. Beads were then pelleted and washed three times with 500 μl of lysis buffer, resuspended in 1/10 original lysate volume of denaturing sample buffer, and processed for immunoblotting and detection.

### Sequence Analysis

In **Figure 1A**, SH3 binding sites in the PRD of Shi are predicted based on the sequence of Shi-L (Shi-PJ) and tools developed in (Kundu et al., 2013; Kundu et al., 2014) found at http://modpepint.informatik.uni-freiburg.de/SH3PepInt/. Actin binding sites in the PRD of Shi-L were predicted and tested in (Zhang et al., 2020). Mammalian dynamin phosphorylation sites in the PRD are predicted based on tools developed in (Hornbeck et al., 2015) found at https://www.phosphosite.org. Shi phosphorylation sites in the PRD are predicted based on the sequence of Shi-L (Hu et al., 2019) found at https://www.flyrnai.org/tools/iproteindb. Calcineurin binding sites were predicted and tested in (Xue et al., 2011).

For conservation studies, NCBI was used to identify organisms with long and short PRD isoforms. The following accession numbers were used. *Anopheles gambiae* XP_061497779 and XP_061497784, *Tribolium castaneum:* XP_064214223.1 and XP_064214222.1 and *Trichonephila clavata:* GFQ72306 and GFQ72301.

### Statistical analysis

Datasets were first analyzed with the D’Agostino-Pearson normality test. Normally distributed datasets were analyzed with either an unpaired two-sided Student’s t-test (for datasets with two groups) or a one-way ANOVA with Tukey’s multiple comparisons (for datasets with more than two groups), while data that is not normally distributed as analyzed with either a two-sided Mann-Whitney U test (two groups) or a Kruskal-Wallis test with Dunn’s multiple comparisons (more than two groups). Error bars report ± s.e.m. For graphs with more than one statistical comparison, no symbol means not significant (ns). Otherwise, ns= p>0.05, * = P < 0.05; ** = P < 0.01; *** = P <0.001. Detailed information about genotypes, sample sizes, and statistical analyses performed for each dataset can be found in **Supplemental Table 3**.

## Acknowledgements

We thank Michael Marr, Steven DeLuca, Erica Dresselhaus, Lindsay Case for technical assistance and/or feedback on the manuscript, Graeme Davis for antibodies, the Bloomington *Drosophila* Stock Center (Indiana University, Bloomington, IN; National Institutes of Health [NIH] P40OD018537) for providing fly stocks, and the DSHB (created by the National Institute of Child Health and Human Development of the NIH) for monoclonal antibodies. We especially thank Greg Hoeprich and Berith Isaac for their guidance and technical assistance with TIRF microscopy and electron microscopy, respectively. We thank the Brandeis University Louise Mashal Gabbay Cellular Visualization Center Electron Microscopy Core Facility (RRID:SCR_026272) and the Brandeis Light Microscopy Core Facility (RRID:SCR_025892) for technical support and access to instruments. This work was supported by grants S10 OD034223 for the Abberior Facility Line STED microscope, R01 NS116375 (A.A.R.), R35 GM134895 (B.L.G.), T32 GM139798 (A.M.S, L.J.W.), and T32 NS2007292 (K.M.D.L.G.).

---

**Supplemental Figure 1:**
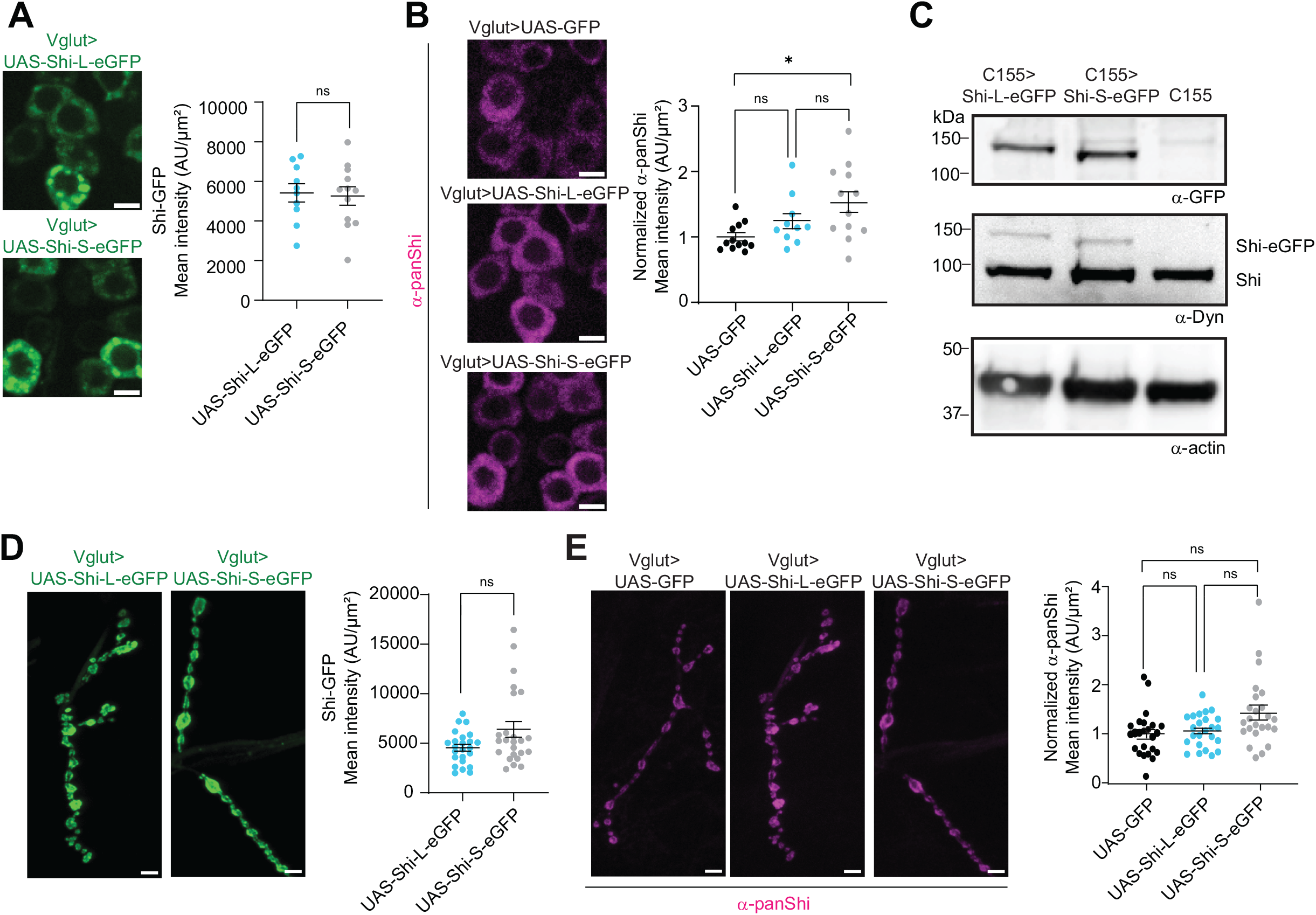
Characterizing Shi isoform overexpression tools. UAS-Shi-L-eGFP and Shi-S-eGFP transgenes were expressed under the control of the pan-neuronal driver GAL4^C155^ or the motor neuron driver GAL4^Vglut^. **(A-B)** Spinning disk MaxIPs of motor neuron cell bodies showing **(A)** Shi-GFP fluorescence or **(B)** total Shi stained with mouse-α-pan Shi. **(C)** Western blot of adult *Drosophila* heads stained with chicken-α-GFP and rabbit-α-pan Shi. **(D-E)** Spinning disk MaxIPs of muscle 4 NMJs overexpressing Shi-L-eGFP and Shi-S-eGFP showing **(D)** Shi-GFP or **(E)** total Shi stained with mouse-α-pan Shi. All scale bars 5µm. Graphs show mean +/− SEM. N = average intensity of cell bodies per larvae **(A-B)** or single NMJ (**D-E)**. See **Supplemental Table 3** for detailed genotypes, sample sizes, and statistical analyses.

**Supplemental Figure 2:**
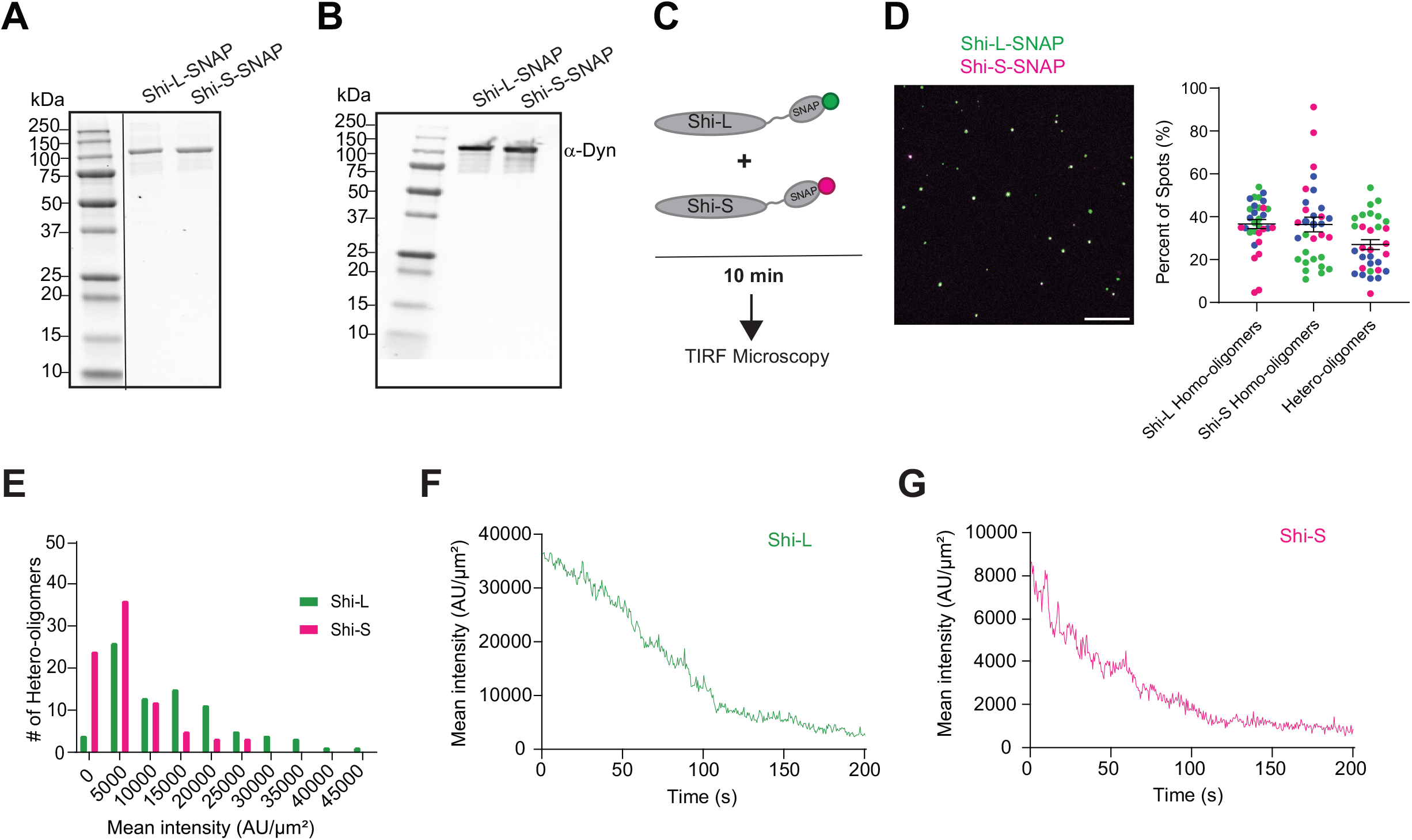
Shi-L and Shi-S can co-oligomerize into higher order structures. **(A)** SDS-PAGE gel stained with Coomassie Blue showing purified Shi proteins isolated from baculovirus-infected Sf9 cells. **(B)** Western blot stained with mouse-α-pan Shi shows purified Shi-L-SNAP and Shi-S-SNAP are the dominant species, and there is minimal contamination by endogenous Sf9 cell dynamin, which is predicted to be recognized by this antibody and would have a lower molecular weight. **(C)** 15 nM each of purified Shi-L-SNAP^549^ (green) and Shi-S-SNAP^488^ (magenta) were mixed and pre-incubated for 10 minutes under low salt conditions (50 mM KCl, 1 mM MgCl_2_), then passively absorbed to the bio-PEG-coated coverslip and directly visualized by TIRF microscopy. Scale bar is 10 μm. **(D)** The percentage of spots that formed hetero-oligomers (two colors) and homo-oligomers (one color) was manually counted in three independent experiments (each experiment represented by different color dots). Graph shows mean +/− sem. **(E)** Histogram showing the starting mean intensities of Shi-L-SNAP^549^ and Shi-S-SNAP^488^ in a hetero-oligomer. Oligomers were then photobleached and intensity was measured over time. Example trace showing the gradual decay in mean intensity of **(F)** Shi-L-SNAP^549^ and **(G)** Shi-S-SNAP^488^ in a hetero-oligomer.

**Supplemental Figure 3:**
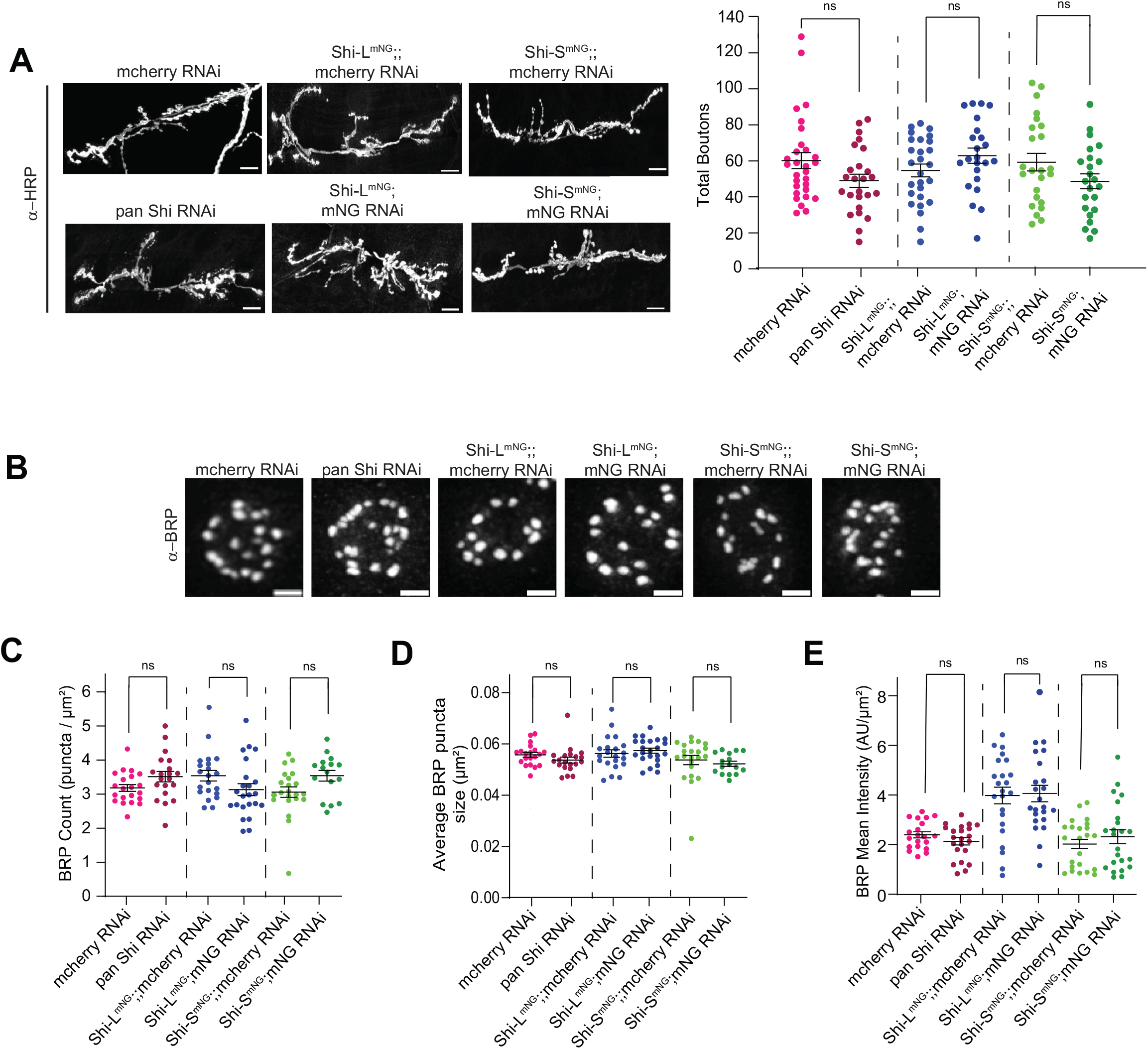
Isoform-specific knockdown of Shi-L^mNG^ and Shi-S^mNG^ at the NMJ does not affect synaptic morphology or active zone distribution. UAS-mNG RNAi or control mcherry RNAi were expressed under the control of the motor neuron driver GAL4^Vglut^. **(A)** Spinning disk Max IPs of muscle 6/7 NMJs in controls and knockdowns stained with α-HRP. HRP intensities are not matched across conditions. Scale bar is 10 µm. **(B)** STED microscopy MaxIPs of muscle 4 NMJs in controls and knockdowns stained with α-BRP to label active zones. Scale bar is 1 μm. **(C**-**E)** Quantification of active zone parameters. Graphs show mean +/− SEM; N=single NMJ. See **Supplemental Table 3** for detailed genotypes, sample sizes, and statistical analyses.

**Supplemental Figure 4:**
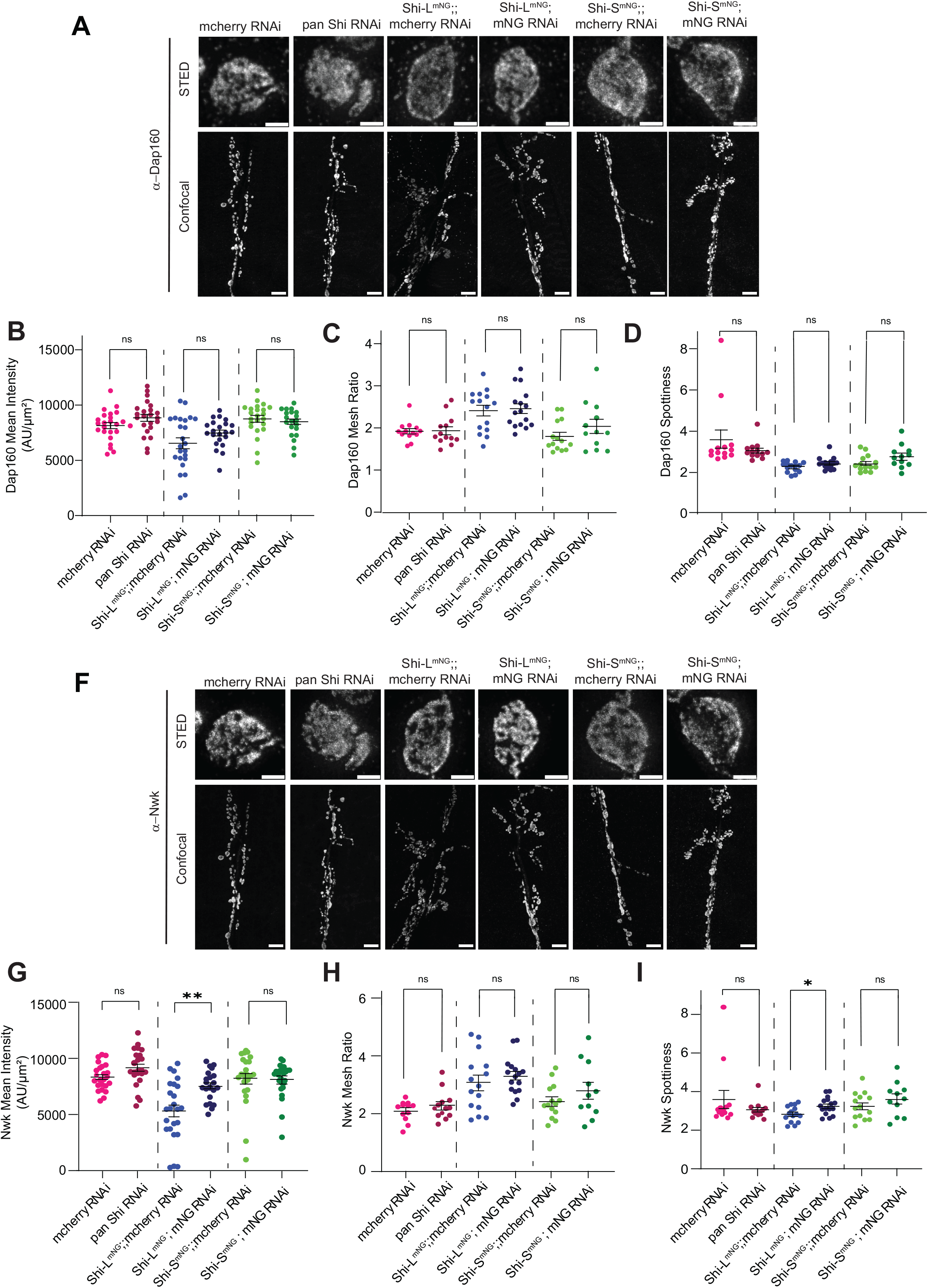
Knockdown of Shi-L^mNG^ or Shi-S^mNG^ does not affect the levels or distribution of periactive zone proteins. **(A)** MaxIPs of muscle 6/7 NMJs in controls and knockdowns stained with α-Dap160. Images were collected on a STED microscope (top, scale bar is 1 µm) and spinning disc confocal microscope (bottom, scale bar is 10 µm). STED images are not intensity matched to each other, though they were collected and analyzed with the same settings. **(B-D)** Quantification of periactive zone protein levels and organization. **(E-H)** Same as A-D but stained with α-Nwk instead of α-Dap160. Note that samples in this figure are all from the same dataset. Graphs show mean +/− SEM. N= single NMJ. See **Supplemental Table 3** for detailed genotypes, sample sizes, and statistical analyses.

**Supplemental Figure 5:**
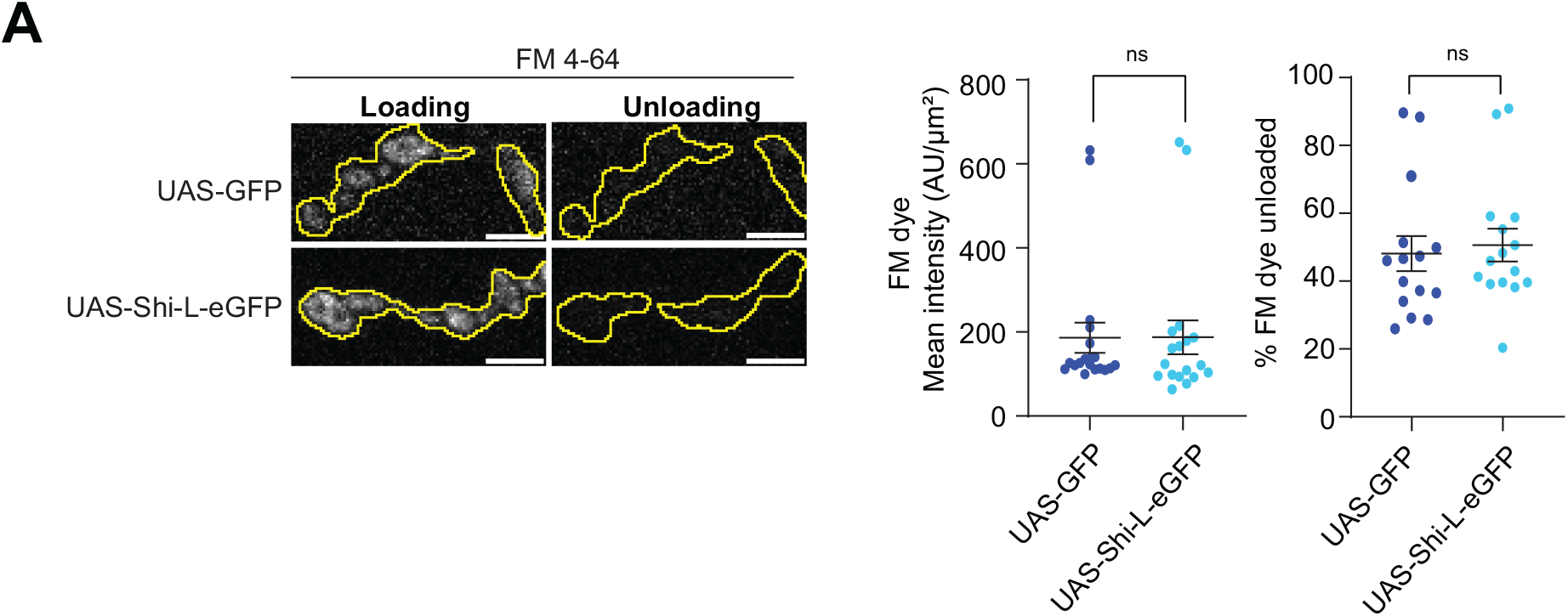
Overexpression of Shi-L-eGFP does not affect FM dye loading or unloading. **(A)** Spinning disk MaxIPs of muscle 6/7 NMJs. NMJs were stimulated with 90 mM KCl and 2 mM CaCl_2_ for 5 minutes to load with FM 4-64 dye (magenta), imaged, stimulated again with 90 mM KCl and 2 mM CaCl_2_ for 2 minutes to unload the dye and then imaged a second time. Scale bar is 5µm. **(B)** Quantification shows mean uptake (FM dye mean intensity +/− SEM) and release (percent FM dye unloaded +/− SEM). N=single NMJ. See **Supplemental Table 3** for detailed genotypes, sample sizes, and statistical analyses.

**Supplemental Figure 6:**
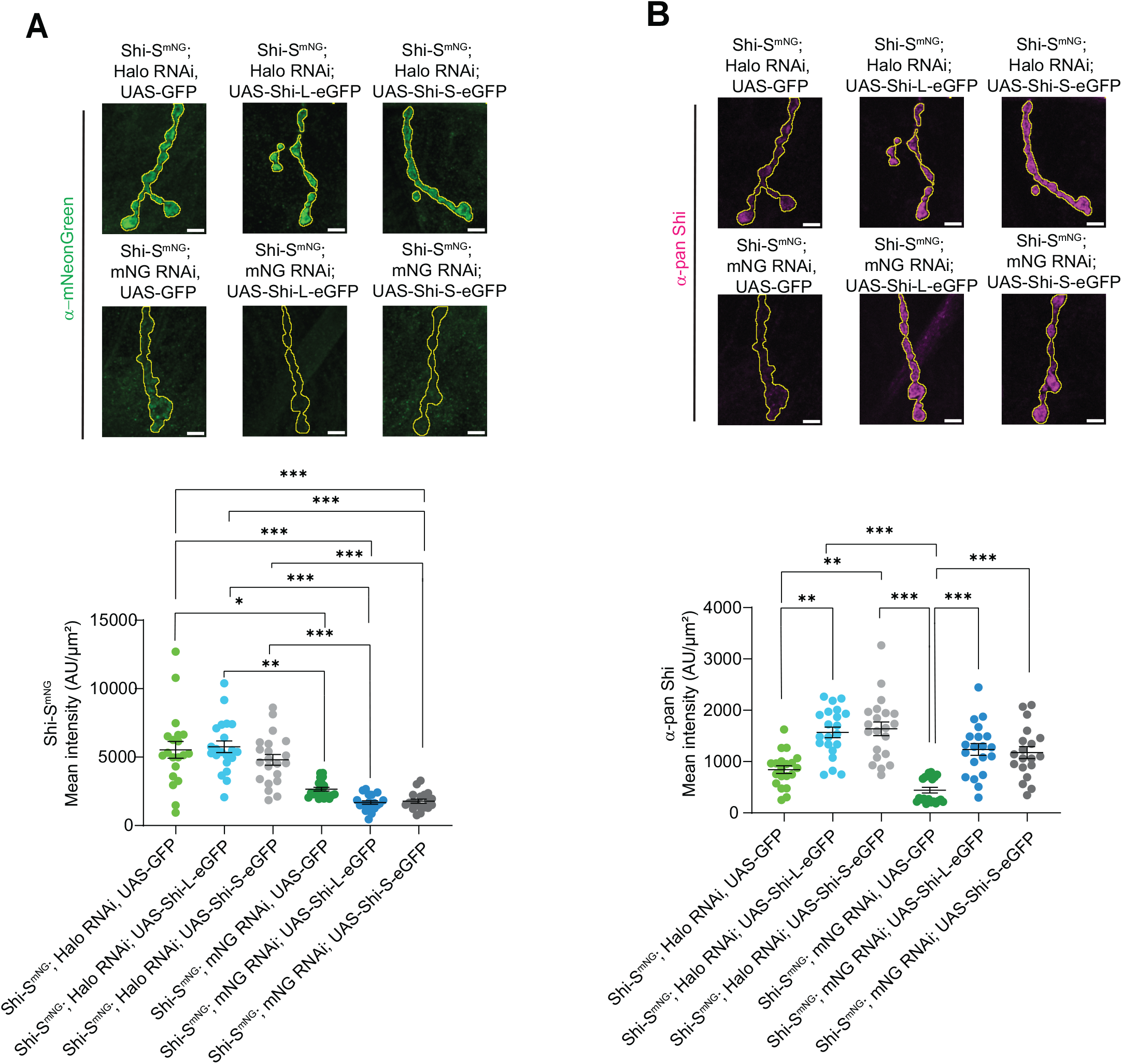
Efficient knockdown and overexpression of Shi isoforms at the NMJ. Spinning disk MaxIPs of muscle 4 NMJs stained with **(A)** chicken-α-mNG or **(B)** mouse-α-pan Shi. Scale bars are 5µm. Graphs show **(A)** knockdown efficiency of Shi-S^mNG^ and **(B)** total Shi levels (mean intensity+/− SEM) when GFP, Shi-L-eGFP or Shi-S-eGFP are co-expressed with an mNGRNAi in motor neurons. See **Supplemental Table 3** for detailed genotypes, sample sizes, and statistical analyses.

**Supplemental Table 1:** Viability of fly stocks and isoform knockdowns.

| Genotype | Survival |
| --- | --- |
| Shi-S<sup>mNG</sup>/y | Viable |
| Shi-L<sup>mNG</sup>/y | Viable |
| Shi-S<sup>Halo</sup>/y | Hemizygous lethal |
| Shi-S<sup>Halo</sup>/y; ;Dp(1:3)Dc133,PBAC[DC133] VK00033/+ | Viable |
| Shi-S<sup>mNG</sup>/y;mNG RNAi/+; tubulin Gal4/+ | Lethal* |
| Shi-L<sup>mNG</sup>/y;mNG RNAi/+; tubulin Gal4/+ | Viable |
| C155 Gal4, Shi-S<sup>mNG</sup>/y; mNG RNAi/+ | Lethal* |
| C155 Gal4, Shi-L<sup>mNG</sup>/y; mNG RNAi/+ | Viable |
| C155 Gal4/y; pan Shi RNAi/+<br>C155 Gal4/+; pan Shi RNAi/+ | Lethal* |

\* No larvae observed

**Supplemental Table 2:** Rescue of lethality of pan neuronal knockdown of Shi-S^mNG^ or both isoforms.

| Genotype * | Observed/expected** |
| --- | --- |
| C155 Gal4, Shi-S<sup>mNG</sup>/y; Halo RNAi, UAS-GFP | 47/34*** |
| C155 Gal4, Shi-S<sup>mNG</sup>/y; mNG RNAi, UAS-GFP/+ | 0/49 |
| C155 Gal4, Shi-S<sup>mNG</sup>/y; Halo RNAi/+; UAS-Shi-S-eGFP/+ | 28/25*** |
| C155 Gal4, Shi-S<sup>mNG</sup>/y; Halo RNAi/+; UAS-Shi-L-eGFP/+ | 42/21*** |
| C155 Gal4, Shi-S<sup>mNG</sup>/y; mNG RNAi/+; UAS-Shi-S-eGFP/+ | 0/26 |
| C155 Gal4, Shi-S<sup>mNG</sup>/y ;mNGRNAi/+; UAS Shi-L eGFP/+ | 0/24 |
| C155 Gal4/y; Halo RNAi, UAS-GFP | 62/98 |
| C155 Gal4/y; pan Shi RNAi, UAS-GFP/+ | 1/122 |
| C155 Gal4/y; Halo RNAi/+; UAS-Shi-S-eGFP/+ | 29/44 |
| C155 Gal4/y; Halo RNAi/+; UAS-Shi-L-eGFP/+ | 50/70 |
| C155 Gal4/y; pan Shi RNAi/+; UAS-Shi-S-eGFP/+ | 27/62 |
| C155 Gal4/y; pan Shi RNAi/+; UAS-Shi-L-eGFP/+ | 0/40 |

\* Raised at 25 degrees
\*\*# of the specified genotype eclosed / # of the specified genotype expected based on total number of flies scored
\*\*\* Overrepresentation of the number of observed flies is likely because some progeny containing balancers have reduced viability

**Supplemental Table 3:** Reporting Table.

| Fig | Value | Genotype/Conditions | N | Statistical Test, P value and Notes |
| --- | --- | --- | --- | --- |
| 1D | Western Blot of Shi levels in fly heads | Shi-L<sup>mNG</sup>/y<br>Shi-S<sup>mNG</sup>/y |  |  |
| 1E | mNG Mean Intensity Brain | Shi-L<sup>mNG</sup>/y<br>Shi-S<sup>mNG</sup>/y | 12 brains/12 larvae<br>13 brains/13 larvae | Unpaired t test<br><0.0001 |
| 1F | mNG Mean Intensity Cell bodies | Shi-L<sup>mNG</sup>/y<br>Shi-S<sup>mNG</sup>/y | 19 cell bodies / 6 larvae<br>29 cell bodies / 7 larvae | Unpaired t test<br><0.001 |
| 1G | mNG Mean Intensity Axons | Shi-L<sup>mNG</sup>/y<br>Shi-S<sup>mNG</sup>/y | 37 axons / 9 larvae<br>39 axons / 9 larvae | Unpaired t test<br>0.001 |
| 1H | mNG Mean Intensity NMJ | Shi-L<sup>mNG</sup>/y<br>Shi-S<sup>mNG</sup>/y | 24 NMJs / 6 larvae<br>23 NMJs / 6 larvae | Unpaired t test<br><0.001 |
| 1I | mNG Mean Intensity Muscle | Shi-L<sup>mNG</sup>/y<br>Shi-S<sup>mNG</sup>/y | 15 muscles/ 4 larvae<br>14 muscles/ 4 larvae | Unpaired t test<br><0.0001 |
|  | # Puncta Muscle | Shi-L<sup>mNG</sup>/y<br>Shi-S<sup>mNG</sup>/y | 24 muscles/ 6 larvae<br>25 muscles/ 6 larvae | Mann-Whitney Test<br>0.0002 |
| S1A | GFP Mean Intensity in cell bodies | Vglut/y; ; UAS-Shi-S-eGFP/+<br>Vglut/y; ; UAS-Shi-L-eGFP/+ | 12 brains/12 larvae<br>10 brains/10 larvae | Unpaired T test<br>0.8160 |
| S1B | α-pan Shi Mean Intensity in cell bodies | Vglut/y; UAS-GFP<br>Vglut/y; ; UAS-Shi-S-eGFP/+<br>Vglut/y; ; UAS-Shi-L-eGFP/+ | 12 brains/12 larvae<br>12 brains/12 larvae<br>10 brains/10 larvae | Kruskal Wallis test<br>0.03 |
| S1C | Western Blot of OE Shi levels in fly heads | C155/y; ; UAS-Shi-S-eGFP/+<br>C155/y; ; UAS-Shi-L-eGFP/+<br>C155/y |  |  |
| S1D | GFP Mean Intensity at NMJ | Vglut/y; ; UAS-Shi-S-eGFP/+<br>Vglut/y; ; UAS-Shi-L-eGFP/+ | 24 NMJs/6 larvae<br>24 NMJs/6 larvae | Mann-Whitney test<br>0.1305 |
| S1E | α-pan Shi Mean Intensity at NMJ | Vglut/y; UAS-GFP<br>Vglut/y; ; UAS-Shi-S-eGFP/+<br>Vglut/y; ; UAS-Shi-L-eGFP/+ | 24 NMJs/6 larvae<br>24 NMJs/6 larvae<br>24 NMJs/6 larvae | Kruskal Wallis test<br>0.06 |
| 2C | Shi Mesh Ratio | Shi-L<sup>mNG</sup>/y<br>Shi-S<sup>mNG</sup>/y | 18 NMJs/7 larvae<br>23 NMJs/7 larvae | Mann-Whitney test<br>0.001 |
| 2D | Shi COV | Shi-L<sup>mNG</sup>/y<br>Shi-S<sup>mNG</sup>/y | 25 NMJs/7 larvae<br>24 NMJs/7 larvae | Unpaired T test<br>0.05 |
| 2E | Shi Spottiness | Shi-L<sup>mNG</sup>/y<br>Shi-S<sup>mNG</sup>/y | 18 NMJs/7 larvae<br>23 NMJs/7 larvae | Mann-Whitney test<br>0.12 |
| 2F | α-pan Shi and Shi<sup>mNG</sup> localization | Shi-L<sup>mNG</sup>/y<br>Shi-S<sup>mNG</sup>/y | 28 NMJs/ 8 larvae<br>26 NMJs/7 larvae |  |
| 2H | Shi-L<sup>mNG</sup> & Shi-S<sup>HaloTag</sup> Pearson's R | Shi-L<sup>mNG</sup>/Shi-S<sup>HaloTag</sup> | 41 NMJs/9 larvae |  |
| 2I | Shi CoV | Shi-L<sup>mNG</sup>/Shi-S<sup>HaloTag</sup> | 41 NMJs/9 larvae | Mann-Whitney test<br><0.0001 |
| 2J | Western Blot of pulldown | Shi-L<sup>mNG</sup>/y |  |  |
| S2D | % of spots | Shi-L-SNAP<sup>549</sup> and Shi-S-SNAP<sup>488</sup> | 31 FOV/3 trials | Kruskal Wallis<br>0.02 |
| S2E | Shi-SNAP Mean Intensity | Shi-L-SNAP<sup>549</sup> and Shi-S-SNAP<sup>488</sup> | 83 oligomers |  |
| 3B | α-pan Shi Mean Intensity | Vglut/y; mNG RNAi/+<br>Vglut/y; pan Shi-RNAi/+ | 23 NMJs/6 larvae<br>20 NMJs/6 larvae | Unpaired T test<br><0.001 |
| 3C | mNG Mean Intensity | Vglut, Shi-L<sup>mNG</sup>/y; ; mcherry RNAi/+<br>Vglut, Shi-L<sup>mNG</sup>/y; mNG RNAi/+ | 16 NMJs/4 larvae<br>16 NMJs/4 larvae | Mann-Whitney test<br><0.001 |
| 3D | mNG Mean Intensity | Vglut, Shi-S<sup>mNG</sup>/y; ; mcherry RNAi/+<br>Vglut, Shi-S<sup>mNG</sup>/y; mNG RNAi/+ | 16 NMJs/4 larvae<br>15 NMJs/4 larvae | Unpaired T test<br><0.001 |
| 3E | α-pan Shi Mean Intensity | Vglut, Shi-L<sup>mNG</sup>/y; ; mcherry RNAi/+<br>Vglut, Shi-L<sup>mNG</sup>/y; mNG RNAi/+<br>Vglut, Shi-S<sup>mNG</sup>/y; ; mcherry RNAi/+<br>Vglut, Shi-S<sup>mNG</sup>/y; mNG RNAi/+ | 21 NMJs/6 larvae<br>21 NMJs/6 larvae<br>24 NMJs/6 larvae<br>22 NMJs/6 larvae |  |
| S3A | # Boutons | Vglut, Shi-L<sup>mNG</sup>/y; ; mcherry RNAi/+<br>Vglut, Shi-L<sup>mNG</sup>/y; mNG RNAi/+ | 27 NMJs/14 larvae<br>24 NMJs/13 larvae | Unpaired T test<br>0.14 |
|  | # Boutons | Vglut, Shi-S<sup>mNG</sup>/y; ; mcherry RNAi/+<br>Vglut, Shi-S<sup>mNG</sup>/y; mNG RNAi/+ | 24 NMJs/13 larvae<br>23 NMJs/14 larvae | Unpaired T test<br>0.10 |
|  | # Boutons | Vglut/y;; mcherry RNAi/+<br>Vglut/y; pan Shi-RNAi/+ | 29 NMJs/16 larvae<br>25 NMJs/15 larvae | Mann-Whitney test<br>0.11 |
| S3C | BRP Count/ NMJ area | Vglut, Shi-L<sup>mNG</sup>/y; ; mcherry RNAi/+<br>Vglut, Shi-L<sup>mNG</sup>/y; mNG RNAi/+ | 21 NMJs/6 larvae<br>23 NMJs/6 larvae | Mann-Whitney test<br>0.0701 |
|  | BRP Count/ NMJ area | Vglut, Shi-S<sup>mNG</sup>/y; ; mcherry RNAi/+<br>Vglut, Shi-S<sup>mNG</sup>/y; mNG RNAi/+ | 22 NMJs/6 larvae<br>16 NMJs/6 larvae | Mann-Whitney test<br>0.0558 |
|  | BRP Count/ NMJ area | Vglut/y; ; mcherry RNAi/+<br>Vglut/y; pan Shi RNAi/+ | 21 NMJs/6 larvae<br>20 NMJs/6 larvae | Unpaired T test<br>0.0742 |
| S3D | Average BRP size | Vglut, Shi-L<sup>mNG</sup>/y; ; mcherry RNAi/+<br>Vglut, Shi-L<sup>mNG</sup>/y; mNG RNAi/+ | 22 NMJs/6 larvae<br>16 NMJs/6 larvae | Unpaired T test<br>0.5073 |
|  | Average BRP size | Vglut, Shi-S<sup>mNG</sup>/y; ; mcherry RNAi/+<br>Vglut, Shi-S<sup>mNG</sup>/y; mNG RNAi/+ | 22 NMJs/6 larvae<br>16 NMJs/6 larvae | Mann-Whitney test<br>0.1428 |
|  | Average BRP size | Vglut/y; ; mcherry RNAi/+<br>Vglut/y; pan Shi RNAi/+ | 20 NMJs/6 larvae<br>20 NMJs/6 larvae | Mann-Whitney test<br>0.0605 |
| S3E | BRP Mean Intensity | Vglut, Shi-L<sup>mNG</sup>/y; ; mcherry RNAi/+<br>Vglut, Shi-L<sup>mNG</sup>/y; mNG RNAi/+ | 23 NMJs/6 larvae<br>24 NMJs/6 larvae | Unpaired T test<br>0.87 |
|  | BRP Mean Intensity | Vglut, Shi-S<sup>mNG</sup>/y; ; mcherry RNAi/+<br>Vglut, Shi-S<sup>mNG</sup>/y; mNG RNAi/+ | 23 NMJs/6 larvae<br>20 NMJs/6 larvae | Unpaired T test<br>0.39 |
|  | BRP Mean Intensity | Vglut/y; ; mcherry RNAi/+<br>Vglut/y; pan Shi RNAi/+ | 20 NMJs/6 larvae<br>23 NMJs/6 larvae | Unpaired T test<br>0.17 |
| S4A | Dap160 Mean Intensity | Vglut, Shi-L<sup>mNG</sup>/y; ; mcherry RNAi/+<br>Vglut, Shi-L<sup>mNG</sup>/y; mNG RNAi/+ | 24 NMJs/6 larvae<br>22 NMJs/6 larvae | Unpaired T test<br>0.1142 |
|  | Dap160 Mean Intensity | Vglut, Shi-S<sup>mNG</sup>/y; ; mcherry RNAi/+<br>Vglut, Shi-S<sup>mNG</sup>/y; mNG RNAi/+ | 23 NMJs/6 larvae<br>24 NMJs/6 larvae | Mann-Whitney test<br>0.3579 |
|  | Dap160 Mean Intensity | Vglut/y; ; mcherry RNAi/+<br>Vglut/y; pan Shi RNAi/+ | 24 NMJs/6 larvae<br>23 NMJs/6 larvae | Unpaired T test<br>0.0982 |
| S4B | Dap160 Mesh Ratio | Vglut, Shi-L<sup>mNG</sup>/y; ; mcherry RNAi/+<br>Vglut, Shi-L<sup>mNG</sup>/y; mNG RNAi/+ | 15 NMJs/4 larvae<br>16 NMJs/4 larvae | Unpaired T test<br>0.79 |
|  | Dap160 Mesh Ratio | Vglut, Shi-S<sup>mNG</sup>/y; ; mcherry RNAi/+<br>Vglut, Shi-S<sup>mNG</sup>/y; mNG RNAi/+ | 14 NMJs/4 larvae<br>12 NMJs/4 larvae | Unpaired T test<br>0.22 |
|  | Dap160<br>Mesh Ratio | Vglut/y; ; mcherry RNAi/+<br>Vglut/y; pan Shi RNAi/+ | 13 NMJs/4 larvae<br>13 NMJs/4 larvae | Mann-Whitney<br>test<br>0.65 |
| S4C | Dap160<br>Spottiness | Vglut, Shi-L<sup>mNG</sup>/y; ; mcherry RNAi/+<br>Vglut, Shi-L<sup>mNG</sup>/y; mNG RNAi/+ | 15 NMJs/4 larvae<br>16 NMJs/4 larvae | Mann-Whitney<br>test<br>0.19 |
|  | Dap160<br>Spottiness | Vglut, Shi-S<sup>mNG</sup>/y; ; mcherry RNAi/+<br>Vglut, Shi-S<sup>mNG</sup>/y; mNG RNAi/+ | 14 NMJs/4 larvae<br>12 NMJs/4 larvae | Unpaired T<br>test<br>0.08 |
|  | Dap160<br>Spottiness | Vglut/y; ; mcherry RNAi/+<br>Vglut/y; pan Shi RNAi/+ | 13 NMJs/4 larvae<br>13 NMJs/4 larvae | Mann-Whitney<br>test<br>0.58 |
| S4G | Nwk Mean<br>Intensity | Vglut, Shi-L<sup>mNG</sup>/y; ; mcherry RNAi/+<br>Vglut, Shi-L<sup>mNG</sup>/y; mNG RNAi/+ | 24 NMJs/6 larvae<br>22 NMJs/6 larvae | Unpaired T<br>test<br>0.0012 |
|  | Nwk Mean<br>Intensity | Vglut, Shi-S<sup>mNG</sup>/y; ; mcherry RNAi/+<br>Vglut, Shi-S<sup>mNG</sup>/y; mNG RNAi/+ | 23 NMJs/6 larvae<br>24 NMJs/6 larvae | Mann-Whitney<br>test<br>0.6502 |
|  | Nwk Mean<br>Intensity | Vglut/y; ; mcherry RNAi/+<br>Vglut/y; pan Shi RNAi/+ | 24 NMJs/6 larvae<br>23 NMJs/6 larvae | Unpaired T<br>test<br>0.0567 |
| S4H | Nwk Mesh<br>Ratio | Vglut, Shi-L<sup>mNG</sup>/y; ; mcherry RNAi/+<br>Vglut, Shi-L<sup>mNG</sup>/y; mNG RNAi/+ | 15 NMJs/4 larvae<br>16 NMJs/4 larvae | Unpaired T<br>test<br>0.4196 |
|  | Nwk Mesh<br>Ratio | Vglut, Shi-S<sup>mNG</sup>/y; ; mcherry RNAi/+<br>Vglut, Shi-S<sup>mNG</sup>/y; mNG RNAi/+ | 14 NMJs/4 larvae<br>12 NMJs/4 larvae | Unpaired T<br>test<br>0.2535 |
|  | Nwk Mesh<br>Ratio | Vglut/y; ; mcherry RNAi/+<br>Vglut/y; pan Shi RNAi/+ | 13 NMJs/4 larvae<br>13 NMJs/4 larvae | Mann-Whitney<br>test<br>0.81 |
|  | Nwk<br>Spottiness | Vglut, Shi-L<sup>mNG</sup>/y; ; mcherry RNAi/+<br>Vglut, Shi-L<sup>mNG</sup>/y; mNG RNAi/+ | 15 NMJs/4 larvae<br>16 NMJs/4 larvae | Unpaired T<br>test<br>0.0112 |
| S4I | Nwk Spottiness | Vglut, Shi-S<sup>mNG</sup>/y; ; mcherry RNAi/+<br>Vglut, Shi-S<sup>mNG</sup>/y; mNG RNAi/+ | 14 NMJs/4 larvae<br>12 NMJs/4 larvae | Unpaired T test<br>0.2595 |
|  | Nwk Spottiness | Vglut/y; ; mcherry RNAi/+<br>Vglut/y; pan Shi RNAi/+ | 13 NMJs/4 larvae<br>13 NMJs/4 larvae | Mann-Whitney test<br>0.5788 |
| 4A | EJP amplitude | Vglut/y; mNG RNAi/+<br>Vglut, Shi-S<sup>mNG</sup>/y; mNG RNAi/+<br>Vglut, Shi-L<sup>mNG</sup>/y; mNG RNAi/+<br>Vglut/y; pan Shi RNAi/+ | 5 NMJs<br>5 NMJs<br>4 NMJs<br>5 NMJs | Kruskal-Wallis test |
| 4C | FM dye Mean Intensity | Vglut/y; ; mcherry RNAi/+<br>Vglut/y; pan Shi RNAi/+ | 17 NMJs/9 larvae<br>22 NMJs/9 larvae | Mann-Whitney test<br>0.17 |
|  | % FM dye unloaded | Vglut/y; ; mcherry RNAi/+<br>Vglut/y; pan Shi RNAi/+ | 17 NMJs/9 larvae<br>22 NMJs/9 larvae | Mann-Whitney test<br>0.0135 |
| 4D | FM dye Mean Intensity | Vglut, Shi-L<sup>mNG</sup>/y; ; mcherry RNAi/+<br>Vglut, Shi-L<sup>mNG</sup>/y; mNG RNAi/+ | 19 NMJs/8 larvae<br>24 NMJs/8 larvae | Unpaired t test<br>0.0709 |
|  | % FM dye unloaded | Vglut, Shi-L<sup>mNG</sup>/y; ; mcherry RNAi/+<br>Vglut, Shi-L<sup>mNG</sup>/y; mNG RNAi/+ | 19 NMJs/8 larvae<br>24 NMJs/8 larvae | Mann-Whitney test<br>0.0906 |
| 4E | FM dye Mean Intensity | Vglut, Shi-S<sup>mNG</sup>/y; ; mcherry RNAi/+<br>Vglut, Shi-S<sup>mNG</sup>/y; mNG RNAi/+ | 28 NMJs/12 larvae<br>36 NMJs/12 larvae | Mann-Whitney test<br><0.0001 |
| 4F | Number of bulk cisterna | REST: Vglut, Shi-S<sup>mNG</sup>/y; ; mcherry RNAi/+<br>REST: Vglut, Shi-S<sup>mNG</sup>/y; mNG RNAi/+<br>STIM: Vglut, Shi-S<sup>mNG</sup>/y; ; mcherry RNAi/+<br>STIM: Vglut, Shi-S<sup>mNG</sup>/y; mNG RNAi/+ | 18 boutons/3 larvae<br>10 boutons/3 larvae<br>10 boutons/3 larvae<br>15 boutons/3 larvae | Two Way ANOVA<br>0.0591 |
| S5B | FM dye<br>Mean<br>Intensity | Vglut/y; UAS-GFP<br>Vglut/y; UAS-Shi-L-eGFP | 21 NMJs/6 larvae<br>19 NMJs/6 larvae | Mann-Whitney<br>test<br>0.4612 |
|  | % FM dye<br>unloaded | Vglut/y; UAS-GFP<br>Vglut/y; UAS-Shi-L-eGFP | 21 NMJs/6 larvae<br>19 NMJs/6 larvae | Unpaired T<br>test<br>0.7248 |
| 5A | EJP<br>amplitude in<br>rescues | Vglut/y; mNG RNAi/+ *<br>Vglut, Shi-S<sup>mNG</sup>/y; mNG RNAi/+ *<br>Vglut, Shi-S<sup>mNG</sup>/y; mNG RNAi/+; UAS-Shi-L-eGFP/+<br>Vglut, Shi-S<sup>mNG</sup>/y; mNG RNAi/+; UAS-Shi-S-eGFP/+ | 5 NMJs<br>5 NMJs<br>5 NMJs<br>4 NMJs | Kruskal-Wallis<br>test<br>*Same as<br>4A |
| S6A | mNG Mean<br>Intensity in<br>rescues | Vglut, Shi-S<sup>mNG</sup>/y; HALO RNAi, UAS-GFP/+<br>Vglut, Shi-S<sup>mNG</sup>/y; HALO RNAi/+; UAS-Shi-L-eGFP/+<br>Vglut, Shi-S<sup>mNG</sup>/y; HALO RNAi/+; UAS-Shi-S-eGFP/+<br>Vglut, Shi-S<sup>mNG</sup>/y; mNG RNAi, UAS-GFP/+<br>Vglut, Shi-S<sup>mNG</sup>/y; mNG RNAi/+; UAS-Shi-L-eGFP/+<br>Vglut, Shi-S<sup>mNG</sup>/y; mNG RNAi/+; UAS-Shi-S-eGFP/+ | 20 NMJs/6 larvae<br>21 NMJs/6 larvae<br>21 NMJs/6 larvae<br>21 NMJs/6 larvae<br>20 NMJs/6 larvae<br>19 NMJs/6 larvae | Kruskal<br>Wallis<br><0.001 |
| S6B | α-pan Shi<br>Mean<br>Intensity<br>in rescues | Vglut, Shi-S<sup>mNG</sup>/y; HALO RNAi, UAS-GFP/+<br>Vglut, Shi-S<sup>mNG</sup>/y; HALO RNAi/+; UAS-Shi-L-eGFP/+<br>Vglut, Shi-S<sup>mNG</sup>/y; HALO RNAi/+; UAS-Shi-S-eGFP/+<br>Vglut, Shi-S<sup>mNG</sup>/y; mNG RNAi, UAS-GFP/+<br>Vglut, Shi-S<sup>mNG</sup>/y; mNG RNAi/+; UAS-Shi-L-eGFP/+<br>Vglut, Shi-S<sup>mNG</sup>/y; mNG RNAi/+; UAS-Shi-S-eGFP/+ | 20 NMJs/6 larvae<br>21 NMJs/6 larvae<br>21 NMJs/6 larvae<br>21 NMJs/6 larvae<br>20 NMJs/6 larvae<br>19 NMJs/6 larvae | Kruskal Wallis<br><0.001 |
| 6B | mEJP<br>Amplitude | Vglut, Shi-S<sup>mNG</sup>/y; HALO RNAi, UAS-GFP/+ | 1893 mEJPs/11 NMJs<br>3022 mEJPs/9 NMJs | Kruskal Wallis<br>test<br>0.005 |
|  |  | Vglut, Shi-S<sup>mNG</sup>/y; mNG RNAi, UAS-GFP/+<br>Vglut, Shi-S<sup>mNG</sup>/y; mNG RNAi/+; UAS-Shi-L-eGFP/+<br>Vglut, Shi-S<sup>mNG</sup>/y; mNG RNAi/+; UAS-Shi-S-eGFP/+ | 1792 mEJPs/10 NMJs<br>1186 mEJPs/10 NMJs |  |
| 6C | mEJP Frequency | Vglut, Shi-S<sup>mNG</sup>/y; HALO RNAi, UAS-GFP/+<br>Vglut, Shi-S<sup>mNG</sup>/y; mNG RNAi, UAS-GFP/+<br>Vglut, Shi-S<sup>mNG</sup>/y; mNG RNAi/+; UAS-Shi-L-eGFP/+<br>Vglut, Shi-S<sup>mNG</sup>/y; mNG RNAi/+; UAS-Shi-S-eGFP/+ | 11 NMJs<br>9 NMJs<br>10 NMJs<br>10 NMJs | One way ANOVA<br><0.001 |
| 6D | EJP amplitude | Vglut, Shi-S<sup>mNG</sup>/y; HALO RNAi, UAS-GFP/+<br>Vglut, Shi-S<sup>mNG</sup>/y; mNG RNAi, UAS-GFP/+<br>Vglut, Shi-S<sup>mNG</sup>/y; mNG RNAi/+; UAS-Shi-L-eGFP/+<br>Vglut, Shi-S<sup>mNG</sup>/y; mNG RNAi/+; UAS-Shi-S-eGFP/+ | 12 NMJs<br>9 NMJs<br>13 NMJs<br>15 NMJs | One way ANOVA<br>0.004 |
| 6E | Quantal Content | Vglut, Shi-S<sup>mNG</sup>/y; HALO RNAi, UAS-GFP/+<br>Vglut, Shi-S<sup>mNG</sup>/y; mNG RNAi, UAS-GFP/+<br>Vglut, Shi-S<sup>mNG</sup>/y; mNG RNAi/+; UAS-Shi-L-eGFP/+<br>Vglut, Shi-S<sup>mNG</sup>/y; mNG RNAi/+; UAS-Shi-S-eGFP/+ | 11 NMJs<br>9 NMJs<br>10 NMJs<br>9 NMJs | One way ANOVA<br><0.001 |

**Supplemental Table 4:** Drosophila Strains.

| Reagent or Resource | Source | Identifier |
| --- | --- | --- |
| C155 Gal4 | (Lin and Goodman, 1994) | FBti0002575 |
| Vglut (X) Gal4 | (Daniels et al., 2008) | FBti0129146 |
| Tubulin-Gal4 | Bloomington *Drosophila* Stock Center | FBtp0002651 |
| *w*<sup>1118</sup> | (Hazelrigg et al., 1984) | FBal0018186 |
| Shi-L<sup>mNG</sup> | This Study |  |
| Shi-S<sup>mNG</sup> | This Study |  |
| Shi-S<sup>HaloTag</sup> | This Study |  |
| UAS-GFP | Leslie Griffith |  |
| UAS-mNeonGreen RNAi | This Study |  |
| UAS-Halo RNAi | This Study |  |
| UAS-VALIUM20-mCherry>attP2 (UAS-mcherry-RNAi) | Bloomington *Drosophila* Stock Center | FBst0035785 |
| UAS-pan Shi-RNAi | VDRC | FBti0116948 |
| UAS-Shi-S-eGFP (PA) (III) | This Study |  |
| UAS-Shi-L-eGFP (PJ) (III) | This Study |  |
| UAS-ShiL-eGFP (II) | This Study |  |

**Supplemental Table 5:** Antibodies.

| Reagent or Resource | Source | Identifier | Concentration * |
| --- | --- | --- | --- |
| Mouse-α-Dyn | BD Biosciences | Clone 41, 610245<br>RRID:AB_397640 | 1:1000 IHC/WB |
| Rabbit-α-Dyn | (Roos and Kelly, 1998) | Ab2074 | 1:1000 IHC<br>1:10,000 WB |
| Mouse-α-actin | DHSB<br>(Lin, 1981) | JLA20<br>RRID: AB_528068 | 1:1000 WB |
| Mouse-α-BRP | DSHB<br>(Wagh et al., 2006) | nc82<br>RRID:AB_2314866 | 1:100 IHC |
| Rabbit-α-Nwk 970 | Coyle et al., 2004 | RRID:AB_2567353 | 1:2000 IHC/WB |
| Rabbit-α-mNeonGreen | Cell Signaling Technologies | 53061 | 1:1000 WB |
| Chicken-α-mNeonGreen | Cancertools | 155277 | 1:500 IHC |
| Mouse-α-mNeonGreen | Chromotek/Proteintech | 32F6 | 1:500 IHC |
| Guinea Pig-α-Dap160 | (Emperador-Melero et al., 2025) | GP14 | 1:1000 IHC/WB |
| Chicken-α-GFP | Aves Labs | Aves GFP-1020 | 1:5000 WB |
| GFP nanobody | Nanotag Biotechnologies | N0304 | 1:500 IHC |
| HRP | Jackson/Fisher |  | 1:250-1:500 IHC |
| 549 HALO | Promega | GA1110 | 500 nM IHC |
| STAR Red | Abberior |  | 1:200 IHC |
| STAR Orange | Abberior |  | 1:200 IHC |
| Alexa 488 | Jackson ImmunoResearch |  | 1:250 IHC<br>1:2000 WB |
| Rhodamine Red-X | Jackson ImmunoResearch |  | 1:500 IHC |
| Alexa 647 | Jackson ImmunoResearch |  | 1:250 IHC |
| Dylight 488 | Rockland |  | 1:5000 WB |
| Dylight 680 | Rockland |  | 1:2000 WB |
| Dylight 800 | Rockland |  | 1:2000 WB |
| SNAP-surface 488 | NEB | S9129S |  |
| SNAP-surface 549 | NEB | S9112S |  |

\*IHC= immunohistochemistry
WB= western blot

**Supplemental Table 6:** Plasmids.

| Reagent or Resource | Source |
| --- | --- |
| pFastBac1 | Thermo Fisher<br>10359016 |
| pFastBac1-Shi-L-SNAP | This Study |
| pFastBac1-Shi-S-SNAP | This Study |

## References

Adams, R., and Bischof, L. (1994). Seeded region growing. IEEE Transactions on Pattern Analysis and Machine Intelligence 16, 641–647.

Afuwape, O.A.T., Chanaday, N.L., Kasap, M., Monteggia, L.M., and Kavalali, E.T. (2025). Persistence of quantal synaptic vesicle recycling in virtual absence of dynamins. J Physiol 603, 6161–6184.

Altschuler, Y., Barbas, S.M., Terlecky, L.J., Tang, K., Hardy, S., Mostov, K.E., and Schmid, S.L. (1998). Redundant and distinct functions for dynamin-1 and dynamin-2 isoforms. J Cell Biol 143, 1871–1881.

Antonny, B., Burd, C., De Camilli, P., Chen, E., Daumke, O., Faelber, K., Ford, M., Frolov, V.A., Frost, A., Hinshaw, J.E., et al. (2016). Membrane fission by dynamin: what we know and what we need to know. EMBO J 35, 2270–2284.

Azarnia Tehran, D., and Maritzen, T. (2022). Endocytic proteins: An expanding repertoire of presynaptic functions. Curr Opin Neurobiol 73, 102519.

Barylko, B., Wang, L., Binns, D.D., Ross, J.A., Tassin, T.C., Collins, K.A., Jameson, D.M., and Albanesi, J.P. (2010). The proline/arginine-rich domain is a major determinant of dynamin self-activation. Biochemistry 49, 10592–10594.

Bhave, M., Mettlen, M., Wang, X., and Schmid, S.L. (2020). Early and nonredundant functions of dynamin isoforms in clathrin-mediated endocytosis. Molecular Biology of the Cell 31, 2035–2047.

Bischof, R.A.a.L. (1994). Seeded Region Growing. IEEE Transactions on Pattern Analysis and Machine Intelligence 16.

Cao, H., Garcia, F., and McNiven, M.A. (1998). Differential distribution of dynamin isoforms in mammalian cells. Mol Biol Cell 9, 2595–2609.

Chanaday, N.L., Cousin, M.A., Milosevic, I., Watanabe, S., and Morgan, J.R. (2019). The Synaptic Vesicle Cycle Revisited: New Insights into the Modes and Mechanisms. J Neurosci 39, 8209–8216.

Chen, M.L., Green, D., Liu, L., Lam, Y.C., Mukai, L., Rao, S., Ramagiri, S., Krishnan, K.S., Engel, J.E., Lin, J.J., et al. (2002). Unique biochemical and behavioral alterations in Drosophila shibire(ts1) mutants imply a conformational state affecting dynamin subcellular distribution and synaptic vesicle cycling. Journal of neurobiology 53, 319–329.

Chen, M.S., Burgess, C.C., Vallee, R.B., and Wadsworth, S.C. (1992). Developmental stage- and tissue-specific expression of shibire, a Drosophila gene involved in endocytosis. J Cell Sci 103 (Pt 3), 619–628.

Chen, M.S., Obar, R.A., Schroeder, C.C., Austin, T.W., Poodry, C.A., Wadsworth, S.C., and Vallee, R.B. (1991). Multiple forms of dynamin are encoded by shibire, a Drosophila gene involved in endocytosis. Nature 351, 583–586.

Cheung, G., and Cousin, M.A. (2019). Synaptic vesicle generation from activity-dependent bulk endosomes requires a dephosphorylation-dependent dynamin-syndapin interaction. J Neurochem 151, 570–583.

Clayton, E.L., Anggono, V., Smillie, K.J., Chau, N., Robinson, P.J., and Cousin, M.A. (2009). The phospho-dependent dynamin-syndapin interaction triggers activity-dependent bulk endocytosis of synaptic vesicles. J Neurosci 29, 7706–7717.

Daniels, R.W., Gelfand, M.V., Collins, C.A., and DiAntonio, A. (2008). Visualizing glutamatergic cell bodies and synapses in Drosophila larval and adult CNS. J Comp Neurol 508, 131–152.

Del Signore, S.J., Mitzner, M.G., Silveira, A.M., Fai, T.G., and Rodal, A.A. (2023). An approach for quantitative mapping of synaptic periactive zone architecture and organization. Molecular Biology of the Cell 34, ar51.

Delgado, R., Maureira, C., Oliva, C., Kidokoro, Y., and Labarca, P. (2000). Size of vesicle pools, rates of mobilization, and recycling at neuromuscular synapses of a Drosophila mutant, shibire. Neuron 28, 941–953.

Dickman, D.K., Horne, J.A., Meinertzhagen, I.A., and Schwarz, T.L. (2005). A slowed classical pathway rather than kiss-and-run mediates endocytosis at synapses lacking synaptojanin and endophilin. Cell 123, 521–533.

Dickman, D.K., Lu, Z., Meinertzhagen, I.A., and Schwarz, T.L. (2006). Altered Synaptic Development and Active Zone Spacing in Endocytosis Mutants. Current Biology 16, 591–598.

Dittman, J., and Ryan, T.A. (2009). Molecular circuitry of endocytosis at nerve terminals. Annual review of cell and developmental biology 25, 133–160.

Emperador-Melero, J., Del Signore, S.J., De León González, K.M., Kaeser, P.S., and Rodal, A.A. (2026). Deployment of endocytic machinery to periactive zones of nerve terminals is independent of active zone assembly and evoked release. Elife 14.

Estes, P.S., Roos, J., van der Bliek, A., Kelly, R.B., Krishnan, K.S., and Ramaswami, M. (1996). Traffic of dynamin within individual Drosophila synaptic boutons relative to compartment-specific markers. The Journal of neuroscience : the official journal of the Society for Neuroscience 16, 5443–5456.

Ferguson, S.M., Brasnjo, G., Hayashi, M., Wölfel, M., Collesi, C., Giovedi, S., Raimondi, A., Gong, L.W., Ariel, P., Paradise, S., et al. (2007). A selective activity-dependent requirement for dynamin 1 in synaptic vesicle endocytosis. Science 316, 570–574.

Gan, Q., and Watanabe, S. (2018). Synaptic Vesicle Endocytosis in Different Model Systems. Front Cell Neurosci 12, 171.

Gass, G.V., Lin, J.J., Scaife, R., and Wu, C.F. (1995). Two isoforms of Drosophila dynamin in wild-type and shibire(ts) neural tissue: different subcellular localization and association mechanisms. J Neurogenet 10, 169–191.

Goel, P., Dufour Bergeron, D., Böhme, M.A., Nunnelly, L., Lehmann, M., Buser, C., Walter, A.M., Sigrist, S.J., and Dickman, D. (2019). Homeostatic scaling of active zone scaffolds maintains global synaptic strength. J Cell Biol 218, 1706–1724.

Gonzalez-Bellido, P.T., Wardill, T.J., Kostyleva, R., Meinertzhagen, I.A., and Juusola, M. (2009). Overexpressing temperature-sensitive dynamin decelerates phototransduction and bundles microtubules in Drosophila photoreceptors. J Neurosci 29, 14199–14210.

Grassart, A., Cheng, A.T., Hong, S.H., Zhang, F., Zenzer, N., Feng, Y., Briner, D.M., Davis, G.D., Malkov, D., and Drubin, D.G. (2014). Actin and dynamin2 dynamics and interplay during clathrin-mediated endocytosis. Journal of Cell Biology 205, 721–735.

Gray, N.W., Fourgeaud, L., Huang, B., Chen, J., Cao, H., Oswald, B.J., Hémar, A., and McNiven, M.A. (2003). Dynamin 3 is a component of the postsynapse, where it interacts with mGluR5 and Homer. Curr Biol 13, 510–515.

Gu, C., Yaddanapudi, S., Weins, A., Osborn, T., Reiser, J., Pollak, M., Hartwig, J., and Sever, S. (2010). Direct dynamin-actin interactions regulate the actin cytoskeleton. Embo j 29, 3593–3606.

Heerssen, H., Fetter, R.D., and Davis, G.W. (2008). Clathrin Dependence of Synaptic-Vesicle Formation at the Drosophila Neuromuscular Junction. Current Biology 18, 401–409.

Hinshaw, J.E., and Schmid, S.L. (1995). Dynamin self-assembles into rings suggesting a mechanism for coated vesicle budding. Nature 374, 190–192.

Hornbeck, P.V., Zhang, B., Murray, B., Kornhauser, J.M., Latham, V., and Skrzypek, E. (2015). PhosphoSitePlus, 2014: mutations, PTMs and recalibrations. Nucleic Acids Res 43, D512–520.

Hu, Y., Sopko, R., Chung, V., Foos, M., Studer, R.A., Landry, S.D., Liu, D., Rabinow, L., Gnad, F., Beltrao, P., et al. (2019). iProteinDB: An Integrative Database of Drosophila Post-translational Modifications. G3 (Bethesda, Md.) 9, 1–11.

Ikeda, K., Ozawa, S., and Hagiwara, S. (1976). Synaptic transmission reversibly conditioned by single-gene mutation in Drosophila melanogaster. Nature 259, 489–491.

Imoto, Y., Raychaudhuri, S., Ma, Y., Fenske, P., Sandoval, E., Itoh, K., Blumrich, E.M., Matsubayashi, H.T., Mamer, L., Zarebidaki, F., et al. (2022). Dynamin is primed at endocytic sites for ultrafast endocytosis. Neuron 110, 2815–2835.e2813.

Imoto, Y., Xue, J., Luo, L., Raychaudhuri, S., Itoh, K., Ma, Y., Craft, G.E., Kwan, A.H., Ogunmowo, T.H., Ho, A., et al. (2024). Dynamin 1xA interacts with Endophilin A1 via its spliced long C-terminus for ultrafast endocytosis. The EMBO Journal 43, 3327–3357.

Jetti, S.K., Crane, A.B., Akbergenova, Y., Aponte-Santiago, N.A., Cunningham, K.L., Whittaker, C.A., and Littleton, J.T. (2023). Molecular logic of synaptic diversity between Drosophila tonic and phasic motoneurons. Neuron 111, 3554–3569.e3557.

Jiang, A., Kudo, K., Gormal, R.S., Ellis, S., Guo, S., Wallis, T.P., Longfield, S.F., Robinson, P.J., Johnson, M.E., Joensuu, M., et al. (2024). Dynamin1 long- and short-tail isoforms exploit distinct recruitment and spatial patterns to form endocytic nanoclusters. Nat Commun 15, 4060.

Kasprowicz, J., Kuenen, S., Miskiewicz, K., Habets, R.L., Smitz, L., and Verstreken, P. (2008). Inactivation of clathrin heavy chain inhibits synaptic recycling but allows bulk membrane uptake. J Cell Biol 182, 1007–1016.

Kasprowicz, J., Kuenen, S., Swerts, J., Miskiewicz, K., and Verstreken, P. (2014). Dynamin photoinactivation blocks Clathrin and α-adaptin recruitment and induces bulk membrane retrieval. J Cell Biol 204, 1141–1156.

Koenig, J.H., and Ikeda, K. (1983). Evidence for a presynaptic blockage of transmission in a temperature-sensitive mutant of Drosophila. Journal of neurobiology 14, 411–419.

Koenig, J.H., and Ikeda, K. (1989). Disappearance and reformation of synaptic vesicle membrane upon transmitter release observed under reversible blockage of membrane retrieval. J Neurosci 9, 3844–3860.

Kokotos, A.C., and Cousin, M.A. (2015). Synaptic vesicle generation from central nerve terminal endosomes. Traffic 16, 229–240.

Kokotos, A.C., and Low, D.W. (2015). Myosin II and Dynamin Control Actin Rings to Mediate Fission during Activity-Dependent Bulk Endocytosis. The Journal of Neuroscience 35, 8687–8688.

Kononenko, Natalia L., and Haucke, V. (2015). Molecular Mechanisms of Presynaptic Membrane Retrieval and Synaptic Vesicle Reformation. Neuron 85, 484–496.

Kononenko, Natalia L., Puchkov, D., Classen, Gala A., Walter, Alexander M., Pechstein, A., Sawade, L., Kaempf, N., Trimbuch, T., Lorenz, D., Rosenmund, C., et al. (2014). Clathrin/AP-2 Mediate Synaptic Vesicle Reformation from Endosome-like Vacuoles but Are Not Essential for Membrane Retrieval at Central Synapses. Neuron 82, 981–988.

Kundu, K., Costa, F., and Backofen, R. (2013). A graph kernel approach for alignment-free domain-peptide interaction prediction with an application to human SH3 domains. Bioinformatics 29, i335–343.

Kundu, K., Mann, M., Costa, F., and Backofen, R. (2014). MoDPepInt: an interactive web server for prediction of modular domain-peptide interactions. Bioinformatics 30, 2668–2669.

Liu, Y.W., Neumann, S., Ramachandran, R., Ferguson, S.M., Pucadyil, T.J., and Schmid, S.L. (2011). Differential curvature sensing and generating activities of dynamin isoforms provide opportunities for tissue-specific regulation. Proc Natl Acad Sci U S A 108, E234–242.

Liu, Y.W., Surka, M.C., Schroeter, T., Lukiyanchuk, V., and Schmid, S.L. (2008). Isoform and splice-variant specific functions of dynamin-2 revealed by analysis of conditional knock-out cells. Mol Biol Cell 19, 5347–5359.

Mahr, A., and Aberle, H. (2006). The expression pattern of the Drosophila vesicular glutamate transporter: a marker protein for motoneurons and glutamatergic centers in the brain. Gene expression patterns : GEP 6, 299–309.

Meinecke, M., Boucrot, E., Camdere, G., Hon, W.C., Mittal, R., and McMahon, H.T. (2013). Cooperative recruitment of dynamin and BIN/amphiphysin/Rvs (BAR) domain-containing proteins leads to GTP-dependent membrane scission. J Biol Chem 288, 6651–6661.

Mettlen, M., Pucadyil, T., Ramachandran, R., and Schmid, S.L. (2009). Dissecting dynamin’s role in clathrin-mediated endocytosis. Biochem Soc Trans 37, 1022–1026.

Newton, A.J., Kirchhausen, T., and Murthy, V.N. (2006). Inhibition of dynamin completely blocks compensatory synaptic vesicle endocytosis. Proceedings of the National Academy of Sciences 103, 17955–17960.

Nguyen, T.H., Maucort, G., Sullivan, R.K., Schenning, M., Lavidis, N.A., McCluskey, A., Robinson, P.J., and Meunier, F.A. (2012). Actin- and dynamin-dependent maturation of bulk endocytosis restores neurotransmission following synaptic depletion. PLoS One 7, e36913.

O’Connor-Giles, K.M., Ho, L.L., and Ganetzky, B. (2008). Nervous wreck interacts with thickveins and the endocytic machinery to attenuate retrograde BMP signaling during synaptic growth. Neuron 58, 507–518.

Okamoto, P.M., Gamby, C., Wells, D., Fallon, J., and Vallee, R.B. (2001). Dynamin isoform-specific interaction with the shank/ProSAP scaffolding proteins of the postsynaptic density and actin cytoskeleton. J Biol Chem 276, 48458–48465.

Okamoto, P.M., Tripet, B., Litowski, J., Hodges, R.S., and Vallee, R.B. (1999). Multiple distinct coiled-coils are involved in dynamin self-assembly. J Biol Chem 274, 10277–10286.

Perkins, L.A., Holderbaum, L., Tao, R., Hu, Y., Sopko, R., McCall, K., Yang-Zhou, D., Flockhart, I., Binari, R., Shim, H.S., et al. (2015). The Transgenic RNAi Project at Harvard Medical School: Resources and Validation. Genetics 201, 843–852.

Poodry, C.A., Hall, L., and Suzuki, D.T. (1973). Developmental properties of Shibire: a pleiotropic mutation affecting larval and adult locomotion and development. Developmental biology 32, 373–386.

Prichard, K.L., O’Brien, N.S., Murcia, S.R., Baker, J.R., and McCluskey, A. (2021). Role of Clathrin and Dynamin in Clathrin Mediated Endocytosis/Synaptic Vesicle Recycling and Implications in Neurological Diseases. Front Cell Neurosci 15, 754110.

Raimondi, A., Ferguson, S.M., Lou, X., Armbruster, M., Paradise, S., Giovedi, S., Messa, M., Kono, N., Takasaki, J., Cappello, V., et al. (2011). Overlapping role of dynamin isoforms in synaptic vesicle endocytosis. Neuron 70, 1100–1114.

Rodal, A.A., Motola-Barnes, R.N., and Littleton, J.T. (2008). Nervous wreck and Cdc42 cooperate to regulate endocytic actin assembly during synaptic growth. J Neurosci 28, 8316–8325.

Rosendale, M., Van, T.N.N., Grillo-Bosch, D., Sposini, S., Claverie, L., Gauthereau, I., Claverol, S., Choquet, D., Sainlos, M., and Perrais, D. (2019). Functional recruitment of dynamin requires multimeric interactions for efficient endocytosis. Nat Commun 10, 4462.

Roux, A., Koster, G., Lenz, M., Sorre, B., Manneville, J.B., Nassoy, P., and Bassereau, P. (2010). Membrane curvature controls dynamin polymerization. Proc Natl Acad Sci U S A 107, 4141–4146.

Sabeva, N., Cho, R.W., Vasin, A., Gonzalez, A., Littleton, J.T., and Bykhovskaia, M. (2017). Complexin Mutants Reveal Partial Segregation between Recycling Pathways That Drive Evoked and Spontaneous Neurotransmission. J Neurosci 37, 383–396.

Saheki, Y., and De Camilli, P. (2012). Synaptic vesicle endocytosis. Cold Spring Harb Perspect Biol 4, a005645.

Sontag, J.M., Fykse, E.M., Ushkaryov, Y., Liu, J.P., Robinson, P.J., and Südhof, T.C. (1994). Differential expression and regulation of multiple dynamins. J Biol Chem 269, 4547–4554.

Soykan, T., Maritzen, T., and Haucke, V. (2016). Modes and mechanisms of synaptic vesicle recycling. Current Opinion in Neurobiology 39, 17–23.

Sprecher, S.G., Müller, M., Kammermeier, L., Miller, D.F., Kaufman, T.C., Reichert, H., and Hirth, F. (2004). Hox gene cross-regulatory interactions in the embryonic brain of Drosophila. Mechanisms of development 121, 527–536.

Staples, R.R., and Ramaswami, M. (1999). Functional analysis of dynamin isoforms in Drosophila melanogaster. J Neurogenet 13, 119–143.

Sundborger, A.C., and Hinshaw, J.E. (2014). Regulating dynamin dynamics during endocytosis. F1000Prime Rep 6, 85.

Tanifuji, S., Funakoshi-Tago, M., Ueda, F., Kasahara, T., and Mochida, S. (2013). Dynamin isoforms decode action potential firing for synaptic vesicle recycling. J Biol Chem 288, 19050–19059.

Taylor, M.J., Lampe, M., and Merrifield, C.J. (2012). A feedback loop between dynamin and actin recruitment during clathrin-mediated endocytosis. PLoS Biol 10, e1001302.

van de Goor, J., Ramaswami, M., and Kelly, R. (1995). Redistribution of synaptic vesicles and their proteins in temperature-sensitive shibire(ts1) mutant Drosophila. Proc Natl Acad Sci U S A 92, 5739–5743.

van der Bliek, A.M., and Meyerowitz, E.M. (1991). Dynamin-like protein encoded by the Drosophila shibire gene associated with vesicular traffic. Nature 351, 411–414.

Verstreken, P., Kjaerulff, O., Lloyd, T.E., Atkinson, R., Zhou, Y., Meinertzhagen, I.A., and Bellen, H.J. (2002). Endophilin mutations block clathrin-mediated endocytosis but not neurotransmitter release. Cell 109, 101–112.

Verstreken, P., Ohyama, T., and Bellen, H.J. (2008). FM 1-43 labeling of synaptic vesicle pools at the Drosophila neuromuscular junction. Methods Mol Biol 440, 349–369.

Warnock, D.E., Baba, T., and Schmid, S.L. (1997). Ubiquitously expressed dynamin-II has a higher intrinsic GTPase activity and a greater propensity for self-assembly than neuronal dynamin-I. Mol Biol Cell 8, 2553–2562.

Watanabe, S. (2025). Synaptic Vesicle Recycling Through the Lens of Ultrafast Endocytosis. Annu Rev Neurosci 48, 297–310.

Watanabe, S., Liu, Q., Davis, M.W., Hollopeter, G., Thomas, N., Jorgensen, N.B., and Jorgensen, E.M. (2013a). Ultrafast endocytosis at Caenorhabditis elegans neuromuscular junctions. eLife 2, e00723.

Watanabe, S., Rost, B.R., Camacho-Pérez, M., Davis, M.W., Söhl-Kielczynski, B., Rosenmund, C., and Jorgensen, E.M. (2013b). Ultrafast endocytosis at mouse hippocampal synapses. Nature 504, 242–247.

Wong, M.Y., Cavolo, S.L., and Levitan, E.S. (2015). Synaptic neuropeptide release by dynamin-dependent partial release from circulating vesicles. Mol Biol Cell 26, 2466–2474.

Wu, L.G., and Chan, C.Y. (2024). Membrane transformations of fusion and budding. Nat Commun 15, 21.

Wu, Y., O’Toole, E.T., Girard, M., Ritter, B., Messa, M., Liu, X., McPherson, P.S., Ferguson, S.M., and De Camilli, P. (2014). A dynamin 1-, dynamin 3- and clathrin-independent pathway of synaptic vesicle recycling mediated by bulk endocytosis. Elife 3, e01621.

Xie, W., Adayev, T., Zhu, H., Wegiel, J., Wieraszko, A., and Hwang, Y.W. (2012). Activity-dependent phosphorylation of dynamin 1 at serine 857. Biochemistry 51, 6786–6796.

Xue, J., Graham, M.E., Novelle, A.E., Sue, N., Gray, N., McNiven, M.A., Smillie, K.J., Cousin, M.A., and Robinson, P.J. (2011). Calcineurin selectively docks with the dynamin Ixb splice variant to regulate activity-dependent bulk endocytosis. J Biol Chem 286, 30295–30303.

Zhang, R., Lee, D.M., Jimah, J.R., Gerassimov, N., Yang, C., Kim, S., Luvsanjav, D., Winkelman, J., Mettlen, M., Abrams, M.E., et al. (2020). Dynamin regulates the dynamics and mechanical strength of the actin cytoskeleton as a multifilament actin-bundling protein. Nat Cell Biol 22, 674–688.

Zhao, M., Maani, N., and Dowling, J.J. (2018). Dynamin 2 (DNM2) as Cause of, and Modifier for, Human Neuromuscular Disease. Neurotherapeutics 15, 966–975.

